# Enhancing pancreatic islet cell survival via NMDAR antagonist-driven ROR1 activation as revealed by phosphoproteomics

**DOI:** 10.64898/2026.11.09.764890

**Authors:** Philip Kirschner, Celine Weippert, Sonja Hartwig, Bengt-Frederik Belgardt, Stefan Lehr, Daniel Eberhard, Hadi Al-Hasani, Eckhard Lammert

## Abstract

NMDA receptors (NMDARs) control function and survival not only of neurons, but also of insulin-producing pancreatic beta cells. Since insufficient insulin secretion and beta cell death are key events during diabetes development, targeting NMDARs offers a therapeutic approach. Specifically, the NMDAR antagonist dextrorphan (DXO) increases glucose-stimulated insulin secretion in clinical trials. Furthermore, treatment with DXO or optimized DXO derivatives prevents pancreatic islet cell death upon diabetogenic insults. Nonetheless, the signaling pathways induced by NMDAR inhibition are poorly understood. Here, we use complementary approaches, phosphoproteomics and tyrosine kinase profiling, to demonstrate that DXO and its derivatives rapidly regulate multiple intracellular signaling clusters, including mTOR and ERK signaling, in pancreatic islets. Notably, mTOR inhibition, but not ERK inhibition, is sufficient to negate DXÓs protective effects. Further, we show that activation of Receptor Tyrosine Kinase-like Orphan Receptor 1 (ROR1) is a universal event upon NMDAR inhibition in islets. Simultaneous knockdown of *Ror1/ROR1* and its paralogue *Ror2/ROR2* compromises the viability of both mouse and human islets, highlighting the ROR1/2 signaling axis as a promising avenue for enhancing beta cell survival.

## Introduction

Diabetes mellitus is a group of metabolic disorders characterized by chronic dysregulation of blood glucose homeostasis, induced by insufficient insulin secretion, impaired insulin action, or both^1^. Diabetes can lead to severe complications including cardiovascular diseases, nephropathy, retinopathy, and neuropathy, increasing all-cause mortality. With an estimated number of 828 million adults living with diabetes, it poses a significant threat to public health and represents a massive macroeconomic challenge^1–4^.

Type 2 diabetes (T2D), the most prevalent form of diabetes, is characterized by the progressive deterioration of pancreatic islet function in the context of insulin resistance, mostly due to obesity and likely influenced by alterations in the islet cell composition^5–9^. While at first compensating for increased insulin demand, beta cells progressively fail to compensate. During later stages of T2D pathogenesis, beta cells undergo beta cell death, secretory failure and trans-/dedifferentiation of beta cells^10–17^. Current therapeutic approaches generally do not prevent the progressive loss or dysfunction of beta cells^18–20^. Thus, there remains a critical shortage of therapeutic strategies that directly preserve beta cell survival and function.

The antitussive prodrug dextromethorphan (DXM) of dextrorphan (DXO), an N-methyl-D-aspartate receptor (NMDAR) antagonist, was identified to augment glucose-stimulated insulin secretion (GSIS) in human individuals with type 2 diabetes in two placebo-controlled randomized clinical trials^21,22^. NMDARs were initially known for their important role in neuronal function and survival^23^ and targeting NMDAR antagonist MK801 selectively to glucagon-like peptide 1 receptor-positive brain neurons effectively reduces body weight^24^. Beyond its well-documented neuronal and insulinotropic effects^25^, DXO as well as MK801 and several blood brain barrier (BBB)-impermeable DXO derivatives, have been reported to possess islet cell-protective properties in multiple preclinical models and in human islets^21,22,26–32^. These findings suggest that NMDAR antagonism may preserve beta cell viability in addition to enhancing insulin secretion and reducing obesity, but the underlying molecular mechanisms remain incompletely understood.

Mechanistically, DXO was found to induce expression of activating transcription factor-4 (ATF4)^29^, which is needed for beta cell identity and function under hyperglycemic conditions^33^, leading to a metabolic shift from the tricarboxylic acid (TCA) cycle towards serine-linked mitochondrial one-carbon metabolism (OCM)^29^. This shift, along with stabilization of nuclear factor erythroid 2-related factor 2 (NRF2), results in increased nicotinamide adenine dinucleotide phosphate hydrogen (NADPH) levels, improving cellular antioxidative defense in islet cells^29,34^. However, simultaneously silencing three genes within the serine-linked OCM was needed to still only partially affect islet cell viability^29^. These findings suggested that OCM activation alone could not fully account for the cytoprotective effects of NMDAR antagonism, pointing to the existence of additional downstream mechanisms.

Here, we used unbiased phosphoproteome profiling of pancreatic islets following NMDAR antagonism to identify a previously unrecognized receptor tyrosine kinase-like orphan receptor 1 (ROR1)-driven signaling pathway downstream of NMDAR that preserves the viability of both mouse and human islets. Since ROR1 was previously reported to be absent from most adult human tissues (including brain, heart, lung and liver tissue) but is expressed in pancreatic islets where its expression is upregulated under insulin resistance^35,36^, the ROR1 signaling axis offers a promising avenue for to more selectively preventing human beta cell death than untargeted NMDAR antagonists.

## Results

### Global phosphoproteomics of pancreatic islets under NMDAR antagonism

To identify signaling pathways and components required for islet cell preservation, pancreatic islets were treated with the NMDAR antagonist DXO and a BBB-impermeable DXO derivative (**Fig. 1a; Supplemental Fig. 1a, b**). These antagonists were used, since they were previously shown to protect mouse as well as human pancreatic islets from cell death^21,28,29^. They differ in their chemical structures, with the BBB-impermeable DXO derivative being larger and more hydrophilic. Consequently, it has limited penetration across the BBB, thereby reducing the risk of central nervous system (CNS) adverse events characteristic of untargeted NMDAR antagonists^28^. After treating islets with 10 µM of the compounds for 15 minutes, islet proteins were extracted, trypsin-digested, and phosphopeptide-enriched (**Fig. 1a**). The resulting phosphopeptides were subsequently analyzed by using mass spectrometry (**Fig. 1a-c**). A total of 23,652 phosphopeptides were detected, corresponding to 4,304 phosphoproteins (**Supplemental Table 1**). After quality gating, 16,331 phosphopeptides were retained for analysis, yielding 15,593 unique amino acid sequences. The resulting phosphoproteome predominantly comprised of serine and threonine residues, which accounted for 99.7% of all identified phosphosites, in line with their established approximately 99-fold overabundance compared to phospho-tyrosine residues^37^. Under DXO treatment, 766 phosphopeptides were upregulated and 618 phosphopeptides differentially downregulated (|FC| > 1.5, adjusted p < 0.05) (**Fig. 1b, Supplemental Fig. 1c**). In contrast, the DXO derivative-treated pancreatic islets exhibited a more limited proteomic response, with 587 upregulated and 400 downregulated phosphopeptides (**Fig. 1c, Supplemental Fig. 1d**), probably reflecting reduced islet penetration of the DXO derivative due to its greater polarity and size compared to DXO. Accordingly, these numbers of significantly regulated phosphopeptides are within the range of those elicited by glucagon like peptide-1 receptor agonists^38^.

**Figure 1:**
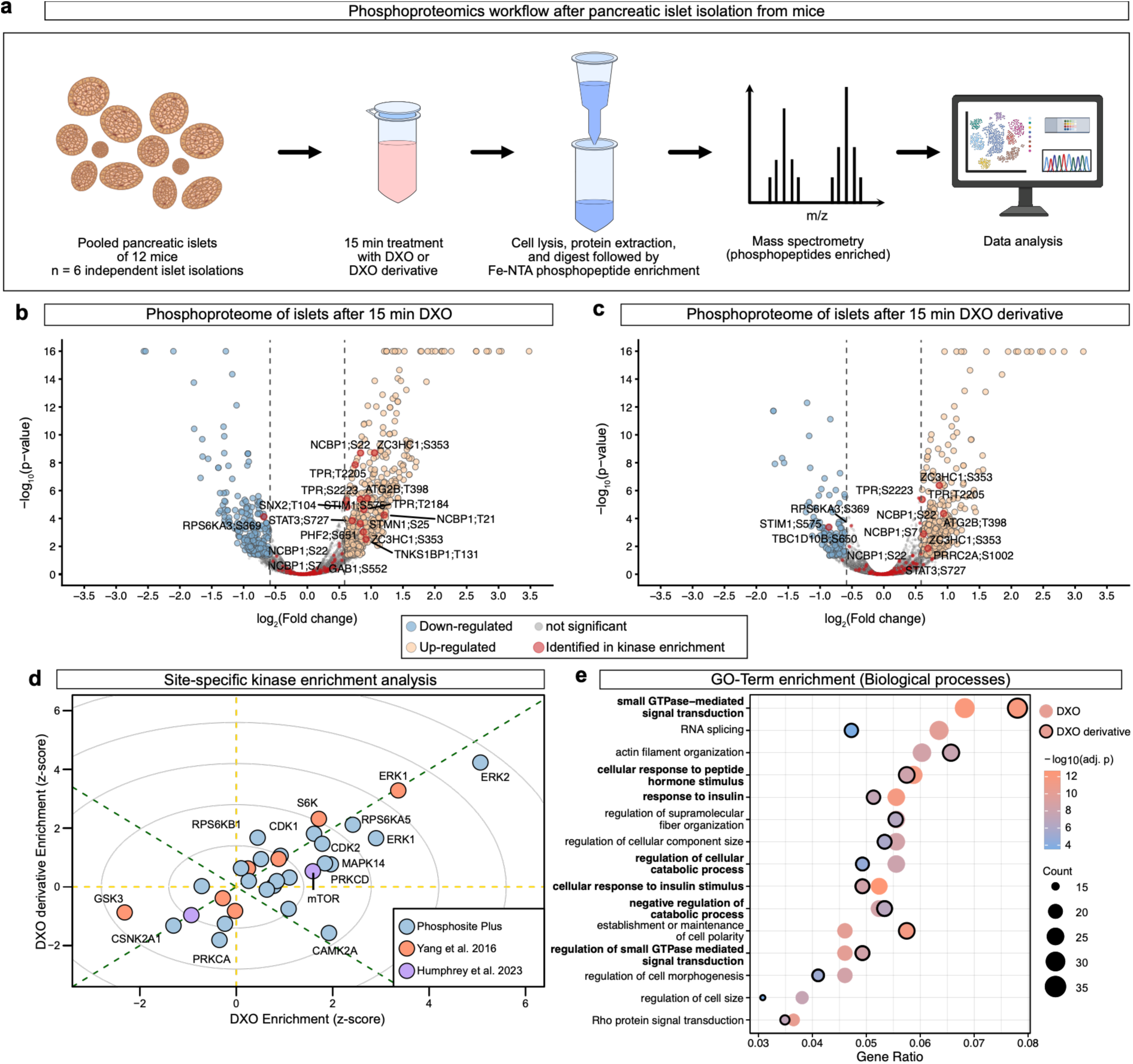
Site-specific kinase enrichment analysis of the phosphoproteome of pancreatic islets treated with NMDA receptor antagonists. **(a)** Schematic illustration of islet treatment with NMDAR antagonist dextrorphan (DXO) or blood brain barrier (BBB)-impermeable DXO derivative, enrichment of phosphopeptides and subsequent mass spectrometry. Data analysis and figure generation were performed in R (4.4.2) using RStudio (2026.09.0+174). **(b, c)** Volcano plots of the islet phosphoproteome after 15 min islet treatment with DXO (b) or DXO derivative^28^ (c) versus vehicle control. Orange dots mark significantly upregulated, blue dots significantly downregulated, and grey dots unchanged phosphopeptides (|FC| > 1.5, as indicated by dashed lines; adj.-p < 0.05). Red dots highlight phosphopeptides associated with signaling pathways identified in downstream kinase enrichment analyses (d) (n = 6 islet preparations). Axis limits were adjusted for visibility. The full data set is presented in supplemental table 1. Uncropped volcano plots are presented in Supplemental Fig. 1c, d. **(d)** Kinase enrichment analysis of the islet phosphoproteome after 15 min islet treatment with DXO and DXO derivative, performed in the RStudio package PhosR^40^ using the perturbPlot2d^39^ function. The x-axis represents the z-score (enrichment score) after DXO treatment, whereas the y-axis represents the z-score after treatment with the DXO derivative. The analysis indicates enhanced or reduced predicted activity of the respective kinases. As PhosR employs different datasets^39,43,125^ for enrichment analysis, these have been colored differently. The dashed green line indicates the line of identity between DXO and DXO derivative enrichment. **(e)** Gene-ontology (GO) term (biological processes) enrichment analysis of significantly changed phosphopeptides (|FC| > 2, adj.-p < 0.05), performed using the clusterProfiler RStudio package. The top 15 GO terms are displayed on the y-axis and were ranked by gene ratio (phosphopeptides in the data set per genes associated with the GO term) on the x-axis (full outputs are shown in supplemental tables 2 and 3). Dot color indicates the Benjamini-Hochberg-adjusted-log10(p-value) (Fisher’s exact test), ranging from p-values of 10^-12^ (orange) to 10^-4^ (blue), and dot size indicates total count of the included phosphopeptides. Results of DXO derivative-treated islets were overlayed and encircled. GO-terms associated with signaling events identified by kinase enrichment (d) are marked in bold on the y-axis.

Site-specific kinase enrichment analysis of the phosphoproteome was performed using the integrated function of PhosR^39,40^ to identify kinases potentially contributing to the signaling induced by DXO or DXO derivative. The analysis indicated activation of kinases associated with the mitogen-activated protein kinase (MAPK, also known as extracellular signal-related kinase (ERK)) signaling pathway, cyclin-dependent kinases (CDKs), mammalian target of rapamycin (mTOR), and the mTOR downstream target ribosomal protein S6 kinase (S6K) beta-1 (**Fig. 1d**). There was also a downregulation of phosphopeptides associated with glycogen synthase kinase-3 beta (GSK3B) and casein kinase 2 subunit alpha 1 (CSNK2A1) by treatment of islets with the NMDAR antagonist DXO (**Fig. 1d, Supplemental Fig. 1e**), in line with reports that GSK3 and casein kinase inhibitors improve insulin secretion and sensitivity^41–43^. Kinase activities upregulated in both DXO-and DXO derivative-treated islets are shown on the green dashed line with the positive slope (i.e., z-scores of 0-6) (**Fig. 1d**). ERK1 stood out the most, with enrichment scores above 4 for both NMDAR antagonists, displaying enrichment in two different reference datasets, while other kinases displayed lower enrichment values (**Fig. 1d**).

To gain further insight into functions of differentially regulated phosphopeptides (|FC| > 2, adjusted p < 0.05), a gene ontology (GO) term enrichment analysis was performed. Significantly regulated phosphopeptides were associated with multiple different signaling events, such as small GTPase-mediated signal transduction and regulation of small GTPase mediated signaling, cellular response to peptide hormone stimulus, negative regulation of catabolic processes, and cellular responses to insulin, indicating that a wide array of anabolic signaling pathways are turned on when NMDARs are inhibited (**Fig. 1e, Supplemental Table 2 and 3**). Further, NMDAR antagonists also triggered phosphorylation events of the cytoskeleton, as indicated by GO-terms “actin filament organization”, “regulation of supramolecular fiber organization”, and “regulation of cell morphogenesis”. Notably, the DXO derivative-treated islets (indicated by outlined dots) displayed comparable effects to those of DXO (indicated by non-outlined dots), closely overlaying the enrichment values of DXO-treated islets (**Fig. 1e, Supplemental Table 3**). Thus, phosphoproteomics analyses led to the identification of ERK and mTOR signaling as well as several biological processes, of which some might be involved in islet cell protection induced by NMDAR inhibition.

### Clustering of phosphopeptide networks indicates alterations in receptor tyrosine kinase signaling and GSIS

We next constructed a functional interaction network of differentially regulated phosphopeptides (|FC| > 2, adjusted p < 0.05) using Search Tool for the Retrieval of Interacting Genes/Proteins (STRING) (**Fig. 2, Supplemental Fig. 2**). To dissect the broader pathways impacted by NMDAR inhibition, the resulting network was segmented via Markov Clustering Algorithm (MCL) (inflation parameter = 1.5; each cluster was assigned a different color), and functional titles were given to these clusters based on enriched gene ontology (GO) terms (**Supplemental Table 4**).

**Figure 2:**
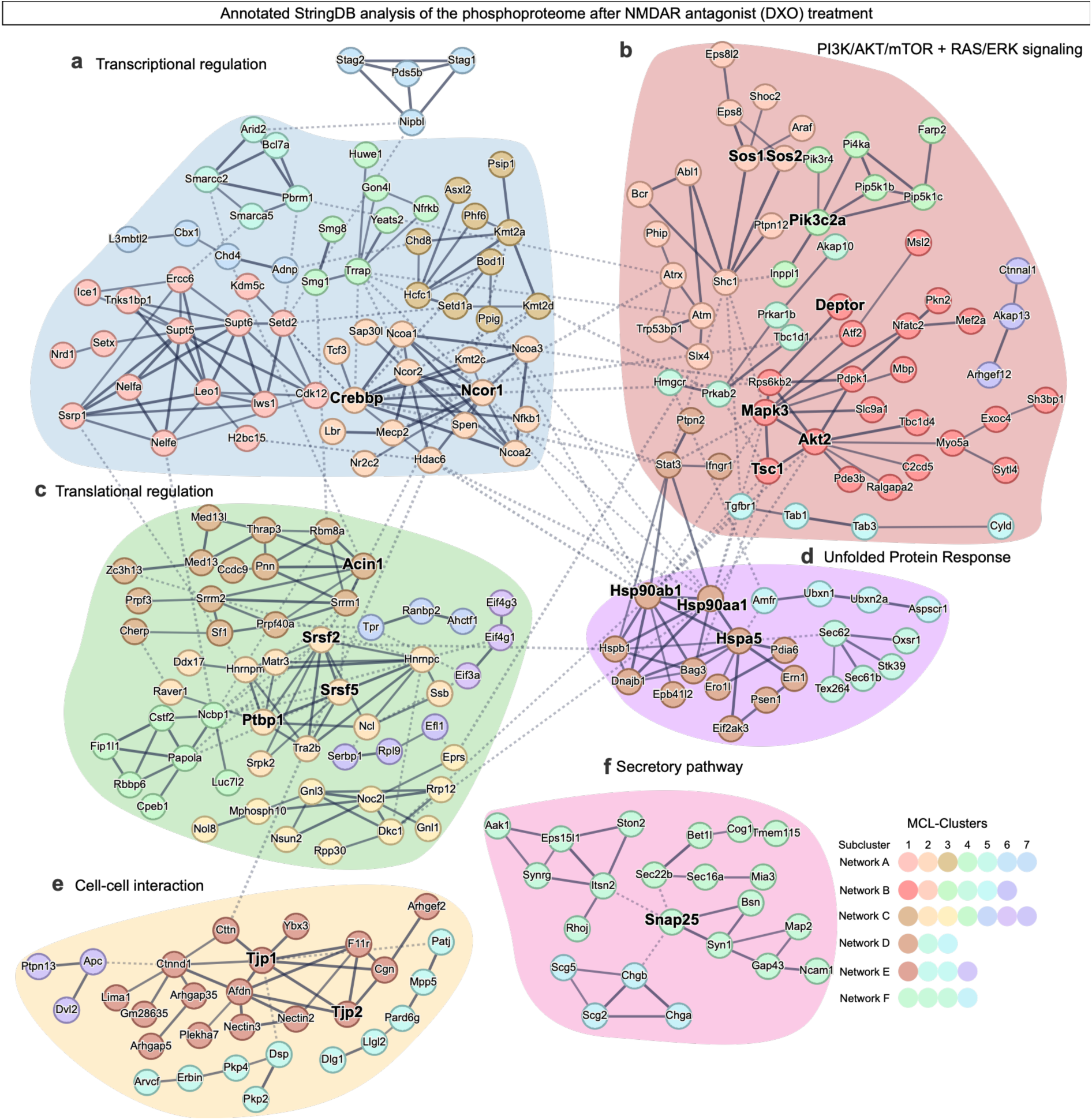
Interaction network of differentially regulated phosphopeptides in response to the treatment of islets with NMDAR receptor antagonist DXO. An annotated protein high-confidence interaction network of significantly altered phosphopeptides in pancreatic islets after 15 min islet treatment with NMDA receptor antagonist DXO (StringDB package in RStudio; interaction score threshold = 750; 306 phosphopeptides total). Disconnected nodes and small networks were omitted for clarity. Connected nodes were partitioned using the Markov clustering algorithm (MCL, inflation parameter = 1.5; each cluster was assigned a color). MCL-clusters were arranged according to the enriched GO-terms of the underlying clusters to gain insights into functional connections, which resulted in the colored fields and respective titles (Table S4): **(a)** Transcriptional regulation (including CREBBP); **(b)** PI3K/AKT/mTOR + RAS/ERK signaling (including PIK3CA2); **(c)** Translational regulation (including PTBP1 and SRSFs); **(d)** Unfolded protein response (including several HSPs); **(e)** Cell-cell interaction (including TJPs); and **(f)** Secretory pathway (including SNAP25).

The largest cluster was prominently associated with transcriptional regulation (**Fig. 2a**), and it encompassed proteins crucial for cell survival, differentiation, secretory function and NMDAR activation (including HDAC/TBL1X/TBLXR1-interacting nuclear receptor co-repressor 1 (NCoR1))^44–47^. Another cluster was linked to peptide hormone signaling, receptor tyrosine kinases (RTKs), and small GTPases, with RAC-alpha serine/threonine-protein kinase 2 (AKT2) – an essential serine/threonine kinase in pancreatic beta cells^48^, serving as a major node (**Fig. 2b**). Notably, our kinase enrichment analysis highlighted mTOR and ERK signaling (**Fig. 1d**), both of which function downstream of multiple RTKs^49,50^. Connected to these pathways was a subcluster associated with phosphatidylinositol processing. This link is noteworthy because phosphoinositide 3-kinase (PI3K) signaling (specifically phosphatidylinositol-4-phosphate 3-kinase C2 domain-containing alpha polypeptide (PIK3C2A), which displayed two significantly upregulated phosphosites in our dataset) has been implicated in shifting autocrine insulin signaling in beta cells toward mitogenic effects^51^ (**Fig. 2b**).

Alterations in translational control and unfolded protein response (UPR) align consistently with our previous RNA sequencing findings and the importance of RNA binding proteins (such as polypyrimidine tract-binding protein 1 (PTBP1) and apoptotic chromatin condensation inducer in the nucleus 1 (ACIN1)) and UPR in beta cells (**Fig. 2c, d**)^29,34,52–56^. The cluster of proteins involved in the UPR also included heat shock protein 90 alpha (HSP90AA1) as a central node protein, which has been recently shown to be involved in survival of alpha cells and beta cells and to link to mTORC1 signaling^57,58^.

A distinct cluster of cell-cell interaction proteins emerged, linking DXO treatment to actin cytoskeletal remodeling and desmosome-associated proteins (**Fig. 2e**), and concurrently, a second cluster related to secretory pathway was identified (**Fig. 2f**), which aligns with the known effects of NMDAR inhibition on GSIS^21,25,28–30^. Although not explicitly shown, structural remodeling of the cytoskeleton likely serves as a functional bridge between these two clusters, since actin remodeling is necessary for physiological GSIS^59^.

Similar clusters emerged in DXO derivative-treated islets (**Supplemental Fig. 2; Supplemental Table 5**). While transcriptional and translational regulation play a major role, the PI3K/AKT/mTOR signaling cluster was smaller compared to DXO-treated islets (**Supplemental Fig. 2a-c**). Nonetheless, PIK3CA2 and associated phosphatidylinositol proteins emerged as nodes in the cluster analysis. Corresponding to the DXO analysis, clusters associated with functions in cell-cell interactions and the secretory pathway also emerged in DXO derivative-treated islets (**Supplemental Fig. 2e**). The BBB-impermeable DXO derivative was designed to be larger and more hydrophilic than the parent compound. Consequently, variations between the two may stem from differences in islet penetration and subsequent time shifts. Overall, profiling of the phosphoproteome reveals that both NMDAR antagonists activate ERK and PI3K/AKT/mTOR signaling.

### NMDAR antagonist DXO requires mTOR activation to preserve pancreatic islets

To validate the kinase enrichment and cluster analysis for ERK and mTOR signaling, we assessed ERK1/2 (MAPK3/1) activation in islets treated with DXO using immunoblotting. Consistent with these analyses, DXO treatment of mouse islets revealed ERK1/2 phosphorylation (T203+Y205/T183+Y185), peaking at 30 to 60 minutes (**Fig. 3a**). Furthermore, 24-hour treatment with increasing concentrations of the NMDAR antagonist induced a dose-dependent increase in P-ERK1/2, reaching a maximum at 10 µM (**Fig. 3b**). Together, these findings demonstrate that DXO induces ERK1/2 phosphorylation in a time-and dose-dependent manner, corroborating our phosphoproteomics data.

**Figure 3:**
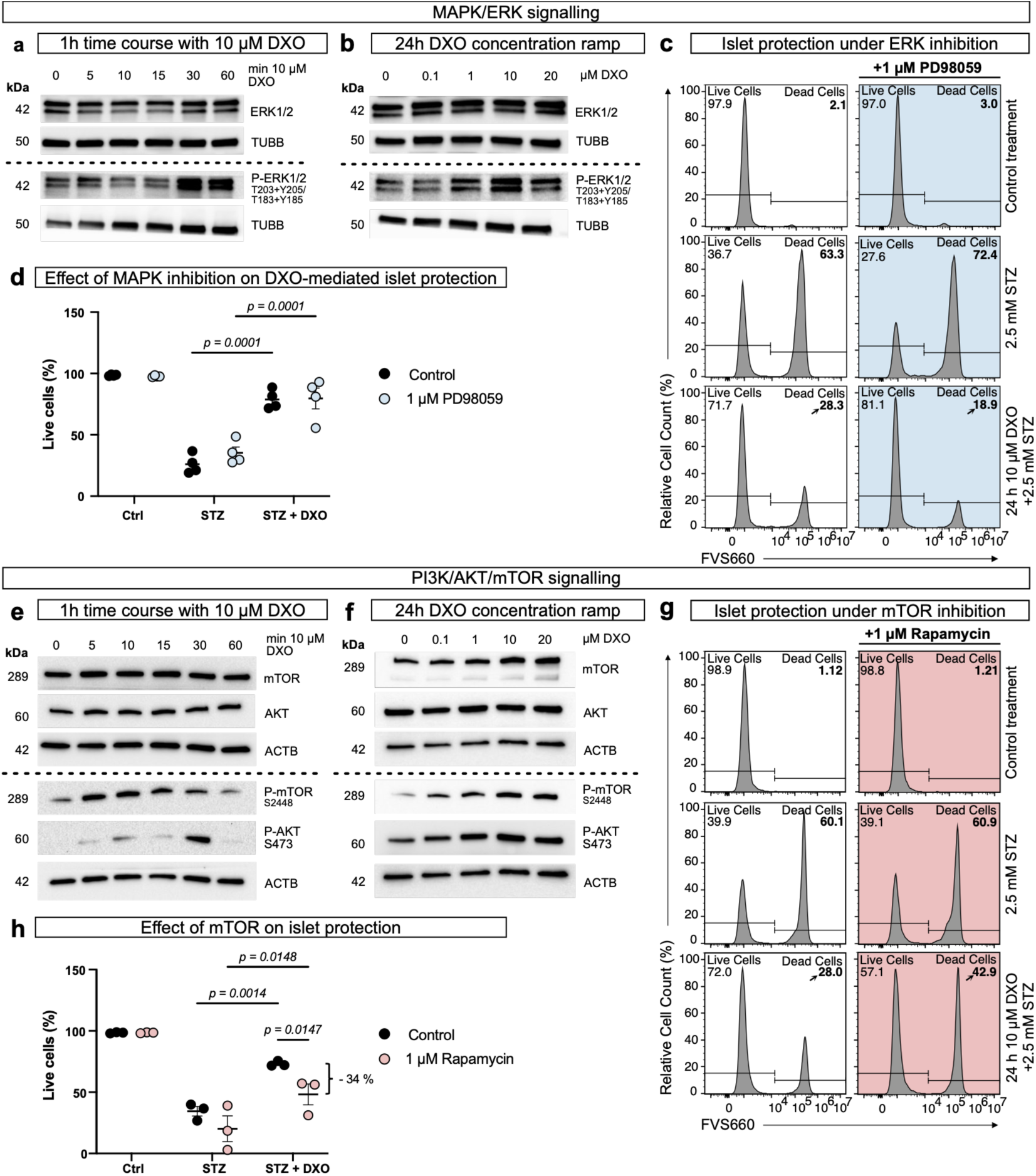
Pancreatic islet cell protection by the NMDA receptor antagonist DXO requires mTOR but not ERK1/2 signaling. **(a)** Time-course immunoblots of DXO treated mouse pancreatic islets. The islets were treated with 10 µM DXO for 0, 5, 10, 15, 30, and 60 minutes. Total ERK1/2 and p-ERK1/2 (T203+Y205/T183+Y185) are shown, and Tubulin beta chain (TUBB) was used as a loading control. The experiment was replicated three times. **(b)** Dose-response immunoblot of mouse pancreatic islets treated for 24h with different concentrations of DXO. Total-ERK1/2 and p-ERK1/2 (T203+Y205/T183+Y185) are shown; TUBB was used as a loading control. The experiment was replicated three times. **(c)** Flow cytometry was used to assess the influence of MEK inhibitor PD98059 (1 µM) on the islet cell-protective effect of DXO. Islets were treated with 10 µM DXO ± inhibitor for 24 h and then treated with 2.5 mM STZ ± DXO ± MEK inhibitor (blue color). Islet cell death was assessed in flow cytometry by fixable viability stain (FVS) 660. **(d)** Percentage of live cells as determined in (c). **(e)** Time-course immunoblots of DXO-treated mouse pancreatic islets. The islets were treated with 10 µM DXO for 0, 5, 10, 15, 30, and 60 minutes. Total mTOR, p-mTOR (S2448), total AKT, and p-AKT(S473) were analyzed; beta actin (ACTB) was used as a loading control. The experiment was replicated three times. **(f)** Dose-response immunoblots of mouse pancreatic islets treated with different concentrations of DXO for 24 h. Total mTOR, p-mTOR (S2448), total-AKT, and p-AKT(S473) were analyzed, and ACTB was used as a loading control. The experiment was replicated five times. **(g)** Flow cytometric analysis of islet cell viability. The role of mTOR inhibitor rapamycin (1µM) was tested in DXO-mediated islet cell protection. Islets were treated with rapamycin for 1 h, before addition of 10 µM DXO and treatment with 2.5 mM STZ ± DXO ± rapamycin for 24 hours. Islet cell death in the presence (red color) or absence of mTOR inhibition was assessed in flow cytometry by FVS660. **(h)** Percentage of living cells determined by flow cytometry analyses as shown in (g). (a, b, e, f) Experiments were repeated 3-5 times. (d, h) A repeated measures two-way ANOVA with Sidak’s post hoc test was performed to test for statistical significance. Flow cytometry data points are presented as mean ± SEM. (c, d, g, h) n = 3-4 independent experiments.

Next, we investigated whether ERK1/2 activation is required for the cytoprotective effect of DXO, since ERK2 has been indicated to be involved in beta cell function and survival using insulinoma cells^60,61^. To inhibit this pathway, we utilized the selective MAPK kinase (MEK) inhibitor PD98059^62^, which reduced DXO-induced ERK1/2 phosphorylation (**Supplemental Fig. 3a**)^62,63^. However, when streptozotocin (STZ; a commonly used toxin to induce beta cell death^64–66^) was used in combination with DXO, the addition of PD98059 failed to abrogate DXO-mediated cytoprotection (**Fig. 3c, d**). These results indicate that while DXO activates ERK1/2, this signaling pathway is not mandatory for its protective effects against STZ-induced cytotoxicity (**Fig. 3c, d**).

We then assessed the roles of mTOR and AKT, as kinase enrichment analyses also identified both as being activated by NMDAR antagonists. Immunoblotting initially validated these predictions: phosphorylation of mTOR (S2448) and AKT (S473) at their kinase-activating sites peaked at 10 and 30 minutes, respectively (**Fig. 3e**). Furthermore, 24-hour treatment with DXO induced the activation of mTOR and AKT in a dose-dependent manner, reaching maximal activation from 1 to 10 µM (**Fig. 3f**). Given that mTOR and AKT operate within the same signaling pathway and that mTOR is required for beta cell survival, insulin secretion and maturation^67–69^, we studied whether mTOR is required for DXO-mediated cytoprotection. Using rapamycin to inhibit mTOR signaling in islets (**Supplemental Fig. 3b**)^70^, we found that disrupting this pathway blocked DXO-mediated protection against STZ-induced cell death (**Fig. 3g, h**). To corroborate this effect, islets treated with two different BBB-impermeable DXO derivatives displayed similar results (**Supplemental Fig. 3c**). Additionally, neither low glucose (5.5 mM) nor S961 (a peptide that selectively inhibits insulin receptor signaling^71^) had any effects on the islet cell-protective effects of DXO (**Supplemental Fig. 4**), demonstrating that a DXO-mediated increase of GSIS (with subsequent insulin receptor activation) is not required for the cell protective effects driven by this NMDAR antagonist.

### Induction of ROR1-associated tyrosine kinase activity by multiple different NMDAR antagonists

We next investigated whether differential tyrosine kinase (TK) activities could be identified in pancreatic islets treated with NMDAR antagonists, since TKs are often upstream of mTOR/AKT signaling events^72,73^. Because phosphotyrosines represented a minority of the total phosphosites enriched by our phosphopeptide enrichment protocol (**Fig. 1**)^37,74^, we utilized the PamGene functional kinase platform to selectively profile these sites, as previously described^75^. More specifically, pancreatic islets were treated with three structurally distinct NMDAR antagonists before their cell lysates were analyzed: DXO, a BBB-impermeable DXO derivative, and MK801, as a highly selective (but not therapeutically used) NMDAR antagonist^76^ (**Fig. 4a** and **Supplemental Fig. 5a-c**). Notably, treatment increased the activity of several TKs that could potentially be upstream of mTOR signaling (**Fig. 4b-e, Supplemental Fig. 5d-f, Supplemental Table 6-8**). Notable targets included multiple src family kinases (SFKs) and EPHA receptors (including EPHA1 and EPHA5), the latter regulating insulin and glucagon secretion from pancreatic islets^77–80^. Strikingly, RTK-like Orphan Receptor-1 (ROR1) was identified as the sole consensus protein with an associated TK activity induced in islets by all three structurally distinct NMDAR antagonists (**Fig. 4e**).

**Figure 4:**
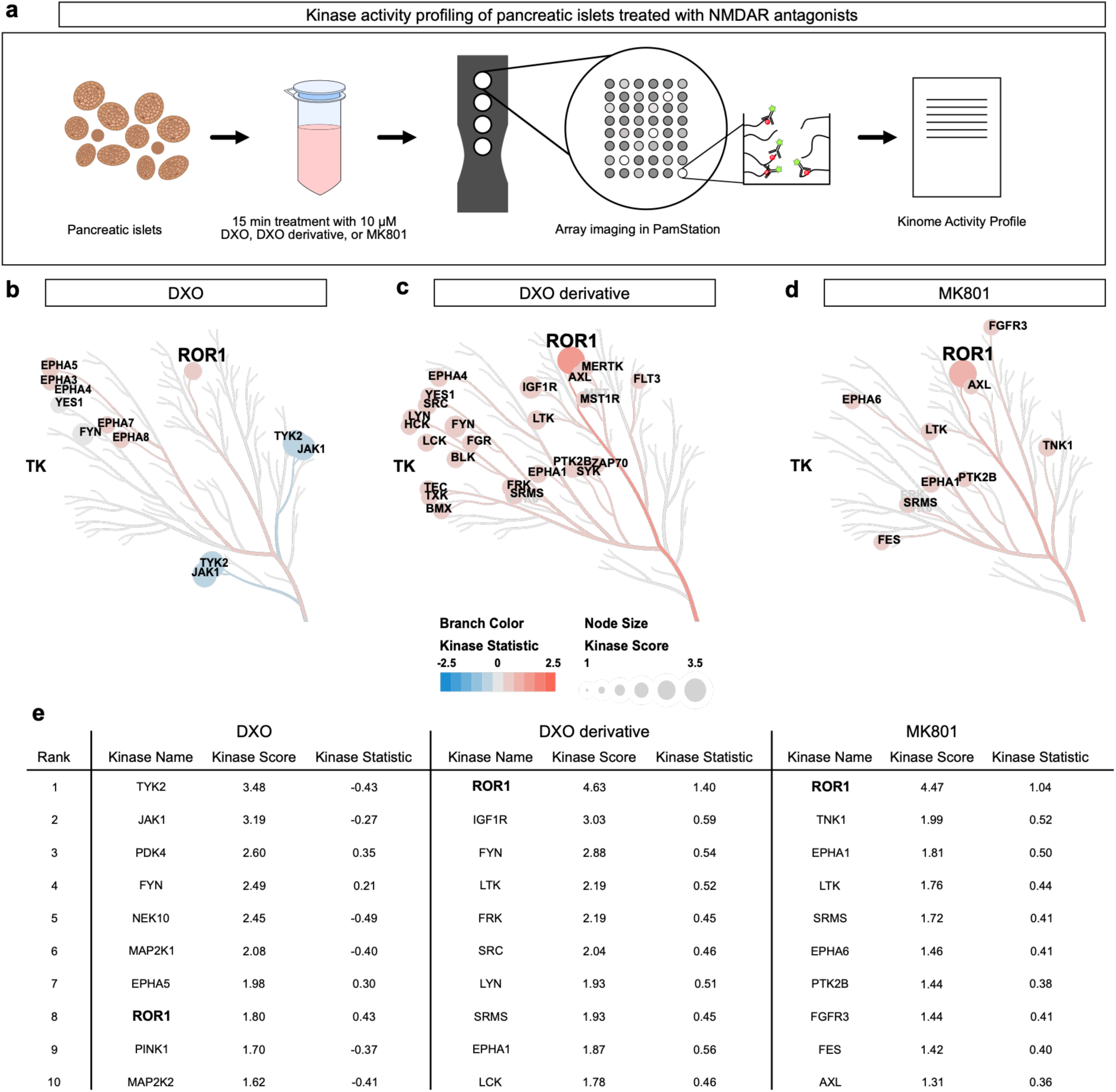
Kinase activity profiling of NMDAR antagonist-treated pancreatic islets unveils ROR1 and EPH receptor tyrosine kinase (RTK) activation. **(a)** Illustration of workflow for islet tyrosine kinome activity profiling via PamGene measurements^126^. Islets were isolated and treated for 15 min with the indicated NMDAR antagonists (n = 6 independent islet isolations). Signal acquisition was performed on the PamStation 12, and raw data were processed by using PamDx B.V. **(b, c, d)** Kinome activity profile visualized on a CORAL kinome tree^127^. Pancreatic islets were treated with 10 µM DXO (b), 10 µM BBB-impermeable DXO derivative^30^ (c), and 10 µM NMDA receptor-selective antagonist MK801^21,76^ (d) for 15 min, and islet protein lysates were subsequently analyzed using the PamGene assay kit. In the CORAL kinome trees, the branch color indicates the “kinase statistic” (up-or downregulation), and the node size depicts the “kinase score” (a measure of significance). **(e)** Table of top 10 kinases for each treatment ranked by “kinase score”. The numeric statistics of the kinome activity profiles for the 10 kinases with the highest kinase score, part of which were depicted in (b-d), ranked by “kinase score” (specificity) with the respective “kinase statistic”, indicating activation or inhibition of the kinases. Full data sets are shown in supplemental tables 6-8. Uncropped CORAL kinome trees are displayed in supplemental fig. 5d-f. TK, tyrosine kinase. Note that kinases that are not canonically listed in the tyrosine kinase family are also not present in the displayed phylogenetic tree (PINK1, PDK4). ROR1 and EPHAs are detected to be activated in the islets treated with either of the three NMDA receptor antagonists.

ROR1 has been shown to be required for normal postnatal development, growth and cell survival, while its protein paralogue, ROR2, appears to possess partially redundant functions and functionally interacts with NMDARs^81–85^. Although predominantly expressed in embryonic rather than adult tissues, ROR1 was reported to be expressed in adult human and primate islets while remaining absent from exocrine pancreatic tissue and most other adult tissues^35^. During tumorigenesis, ROR1 expression is upregulated in tumor cells^81^, consistent with a pro-survival function. Consequently, ROR1 represented a compelling upstream protein candidate activated by NMDAR inhibition.

### ROR1 and ROR2 are needed for full DXO-mediated islet cell preservation

Given that all three islet cell-protective and structurally different NMDAR antagonists enhanced ROR1 activity, we next determined its functional contribution to islet cell survival. Thus, we knocked down *Ror1* in mouse islets before treating them with STZ to induce islet cell death (**Supplemental Fig. 6**). Even though a 90% knockdown efficiency was achieved on mRNA level, islet cell survival was unaltered, neither under control conditions nor STZ treatment with and without DXO. However, we noted a substantial upregulation of its paralogue *Ror2* in islets upon silencing *Ror1* (**Supplemental Fig. 6**), suggesting a functional compensation of ROR1 by ROR2. The latter has been previously reported to be involved in skeletal and cardiac embryonic development^82^. To circumvent the potential issue of functional redundancy, both genes (*Ror1* and *Ror2*) were simultaneously silenced in mouse islets (**Fig. 5a**). After the knockdown had been established on mRNA and protein level (**Fig. 5b, c**), cell viability assays were conducted. These assays revealed that the gene silencing lowered mouse islet cell viability and reduced the protective effect of the NMDAR antagonist DXO (**Fig. 5d, e** and **Supplemental Fig. 6**).

**Figure 5:**
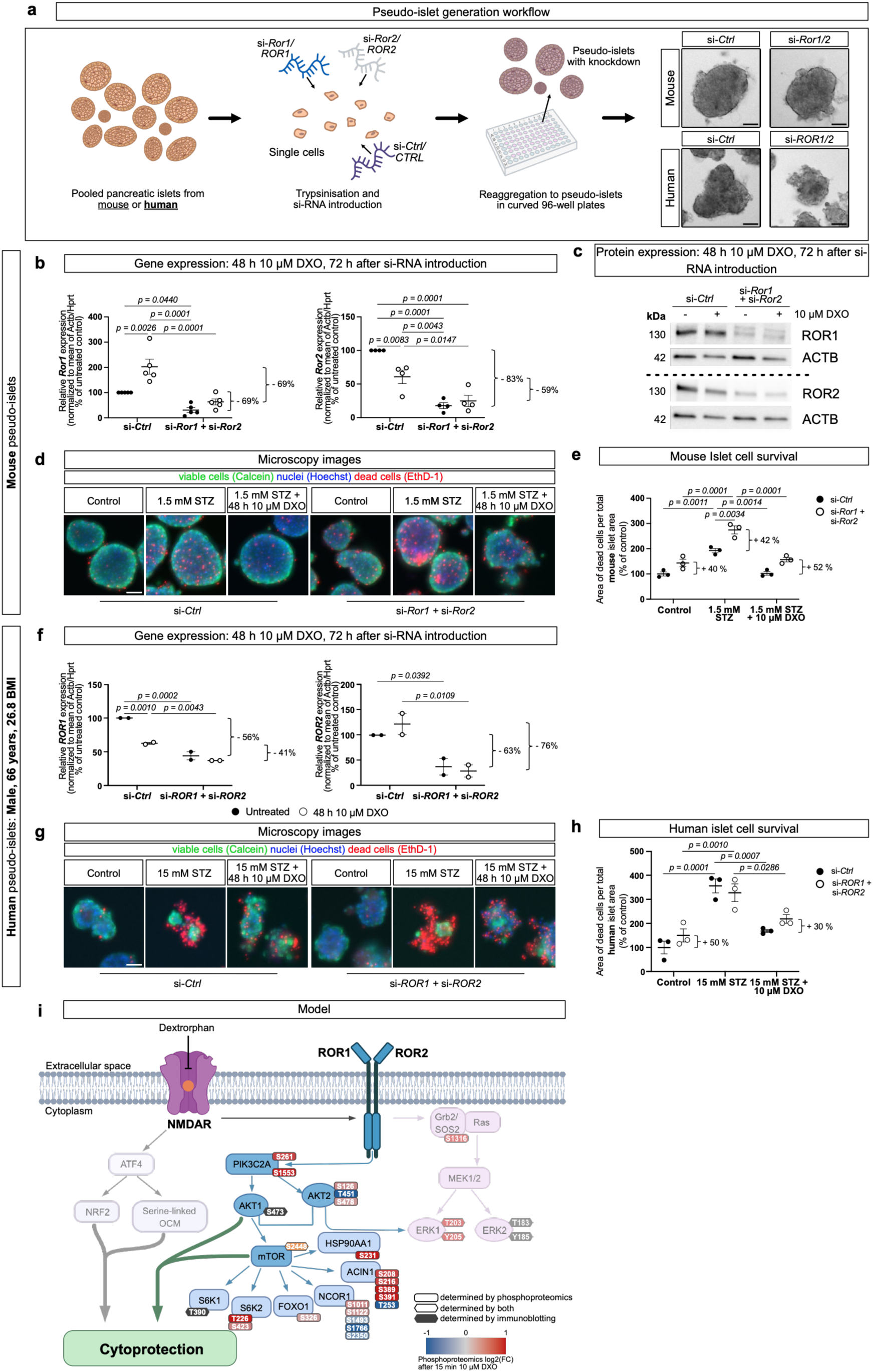
Full pancreatic islet cell survival depends on ROR1/ROR2 expression. **(a)** Illustration of the pseudo-islet generation workflow. Mouse or human pancreatic islets were dispersed, transfected with si-RNA and reaggregated in conical 96-well plates. Mouse and human islets are shown; scale bars: 50 μm. **(b)** Relative mRNA expression of *Ror1* and *Ror2* in si-*Ror1* and si-*Ror2* siRNA transfected mouse pseudo-islets treated with or without 10 µM DXO (48 h). Data are shown as individual values normalized to the housekeeping mRNA *Actb* and *Hprt* relative to si-*Ctrl* siRNA (n = 4-5 individual experiments). **(c)** ROR1 and ROR2 protein expression assessed via immunoblot. ACTB was used as a loading control. Pseudo-islets were transfected with si-*Ctrl* or si-*Ror1* + si-*Ror2* and treated with or without 10 µM DXO (48 h). **(d)** Images of live-dead imaging of si-*Ctrl* versus si-*Ror1* + si-*Ror2* siRNA transfected mouse pseudo-islets treated with or without 1.5 mM STZ (24 h) ± 10 μM DXO (48 h). Nuclei were stained with Hoechst, viable cells with calcein, and dead cells with ethidium-homodimer-1 (Ethd-1) (colors have been adjusted for visibility, scale bar: 50 µm). **(e)** Quantification of the area of dead cells of the experiment shown in (d) (n = 3 stainings). **(f)** Relative mRNA expression of *ROR1* and *ROR2* in si-*ROR1* and si-*ROR2* siRNA transfected human pseudo-islets treated with or without 10 µM DXO (48 h). Data are shown as individual values normalized to the housekeeping mRNA *ACTB* and *HPRT* relative to si-*CTRL* siRNA (n = 2 technical replicates). **(g)** Images of live-dead imaging of si-*CTRL* vs si-*ROR1* + si-*ROR2* siRNA transfected human pseudo-islets treated with or without 1.5 mM STZ (24 h) ± 10 μM DXO (48 h). Nuclei were stained with Hoechst, viable cells with calcein, and dead cells with ethidium-homodimer-1 (Ethd-1). **(h)** Quantification of the area of dead cells of the experiment shown in (g) (n = 3 stainings). **(i)** Illustration of an islet cell-protective pathway derived from our data. NMDAR inhibition (e.g. by dextrorphan, DXO) results in multiple alterations of phosphorylation sites in proteins involved in cell survival, including phospho-sites with a yet unknown function. DXO increases ROR1-associated tyrosine kinase activity, which induces mTOR-signaling dependent cytoprotective effects. Downstream targets such as S6K1, S6K2, FOXO1, NCOR1, ACIN1, and HSP90AA1 displayed up-or downregulated phosphopeptides in phosphoproteomics of 15 min 10 µM DXO. Each phosphosite is colored by its log2 fold change on a diverging scale: blue for decreased and red for increased phosphorylation. To keep smaller changes visible, the color scale was capped at |log2 FC| = 1, All phosphosites quantified with a |log2 FC| > 0.25 on the depicted proteins are shown, regardless of statistical significance. Phosphorylated proteins probed by immunoblotting are indicated by hexagonal boxes. (b, e, f, h) A repeated measures two-way ANOVA with Sidak’s post hoc test was performed to test for statistical significance. Outlier identified in (b) with Grubb’s outlier test (alpha=0.05). Data points are presented as mean ± SEM.

Next, we performed a double knockdown of *ROR1* and *ROR2* in islets from a female and a male human donor (**Fig. 5a** and **Fig. 5f**). STZ concentrations were increased to 15 mM, since human beta cells express the major STZ-transporter (solute carrier family 2, facilitated glucose transporter member 2 (*SLC2A2*/GLUT2)) at lower levels compared to mouse beta cells^86,87^. Silencing both genes in human pseudo-islets recapitulated the phenotype observed in mouse islets to a large extent, in that cell viability was reduced compared to transfection with control siRNA in particular after DXO treatment (**Fig. 5g, h, Supplemental Fig. 6e**). Altogether, these findings identify ROR1 and ROR2 as functionally redundant proteins required for islet cell survival, especially in the context of NMDAR inhibition (**Fig. 5i**).

## Discussion

NMDAR antagonism increases GSIS in mice and humans while effectively halting islet cell death^21,22,27,29,30,32,34^. However, the underlying molecular mechanisms driving this islet cell protection remain largely unknown. Beyond pancreatic effects, NMDAR antagonists have recently emerged as targets for body weight reduction^24^, and they were previously evaluated for neuroprotection in neurodegenerative diseases and stroke^88,89^.

First, a mass spectrometry-based phosphoproteome analysis identified numerous differentially regulated phosphopeptides and novel phospho-sites, establishing a valuable resource for future studies targeting NMDAR in islets and other tissues. Kinase enrichment analyses revealed that both ERK, AKT and mTOR were activated upon NMDAR inhibition indicative of the induction of an immature and more protected beta cell population^8,90^. In beta cells, mTOR signaling is traditionally implicated in growth, proliferation^91–94^, and metabolic regulation, serving as a central hub linked to diverse pathways such as UPR and ATF4 signaling^92,95–98^. Additionally, AKT has been demonstrated to induce beta cell proliferation and protect pancreatic islets and beta cells from diabetogenic stress^99–101^. Previous work showing that the NMDAR antagonist MK801 induces mTOR/p70S6K activation in the rat frontal cortex^102^ points to a conserved therapeutic mechanism across different tissues.

Second, we employed PamGene assay tyrosine kinase profiling, as previously described^75^. This approach identified activation of multiple TKs. Most prominently, NMDAR inhibition activated SFKs, EPHAs, and ROR1. While NMDARs are increasingly recognized to interact with a broad array of proteins, including SFKs, EPHB receptors, and GLURs^103–107^, we have added ROR1. ROR1 has a kinase domain comparable to Trk and insulin receptors and is frequently overexpressed in cancer^84,108–112^.

Here, we propose that NMDAR antagonism activates ROR1, which in combination with the mTOR pathway and other signaling nodes contributes to cytoprotection (**Fig. 5i**). Whereas ROR2 has been shown to interact with NMDARs, ROR1 has not previously been linked to these ion channels^85,113,114^. Since ROR1 and ROR2 are more highly expressed in adult human islets than in most other adult tissues^35,64,115^, targeting ROR1/2 could be more selective than targeting NMDARs, which are more broadly expressed. Thus, besides providing a comprehensive resource for future studies on NMDARs and cell survival, our complementary approaches reveal an islet-specific signaling pathway that could be exploited for diabetes intervention.

## Resource Availiability

### Lead Contact

Further information and request for resources and reagents should be directed to and will be fulfilled by the lead contact Dr. Eckhard Lammert.

### Materials Availability

This study did not generate new unique reagents.

### Data Availability

Source data are provided with this paper. Any raw data supporting this study are available from the corresponding authors. Mass spectrometry proteomics data generated in this study will be deposited to the ProteomeXchange Consortium via the PRIDE^116^ partner repository.

## Acknowledgments

This work was supported and funded by the Heinrich Heine University Düsseldorf, the German Center for Diabetes Research (DZD e.V.), Deutsche Forschungsgemeinschaft (DFG): RTG 2576 (vivid – In vivo investigations towards the early development of type 2 diabetes), CRC 1774 (Cardio-Diabetes-Crosstalk) as well as the Federal Ministry of Health, and the Ministry of Culture and Science of North Rhine-Westphalia. Human islets were provided by the NIDDK-funded Integrated Islet Distribution Program (IIDP; RRID:SCR_014387) at City of Hope, Duarte, CA, USA (NIH Grant U24DK098085). We thank Dr. Michael Turewicz, Maike Müller, Elena Huß, Dr. Roberto Diaz and Dr. Ana Isabel Guzmán Carrasco for insightful discussions and valuable feedback. We would like to acknowledge Barbara Bartosinska, Andrea Köster, and Silke Jakob for technical support.

## Author contribution

P.K. and C.W. performed most of the experiments, with P.K. performing the majority of the studies. S.L., S.H. and H.A. performed the mass spectrometry and phosphoproteomics. B.B. gave advice regarding phosphoproteomics, and D.E. controlled all raw data, checked statistical validity and completeness of data. P.K. generated all Figures with the input from C.W. and E.L.; E.L. guided P.K. and C.W. during their experiments and wrote the manuscript with P.K. All authors read and revised the manuscript and Figures.

## Declaration of generative AI and AI-assisted technologies in the writing process

During the preparation of this work the authors used OpenAI (2023) *ChatGPT* and *Google Gemini* to improve language and readability of the text. After using these tools, the authors reviewed and edited the content as needed and took full responsibility for the content of the published article.

## Methods and Materials

### Mouse model

Male C57BL/6J mice (Janvier), age 9–12 weeks, were used as the source of pancreatic islets for this study. The animals were housed at the Zentrale Einrichtung für Tierforschung und wissenschaftliche Tierschutzaufgaben (ZETT) animal facility at Heinrich-Heine-University Düsseldorf (HHU) under controlled environmental conditions (22 °C, 55% relative humidity, and a 12 h light/12 h dark cycle). Standard laboratory chow and drinking water were provided *ad libitum*. The mice were euthanized by exposure to carbon dioxide (CO₂), followed by cervical dislocation. All procedures involving animal tissues were conducted in accordance with applicable German animal welfare legislation and were approved by the competent authority of the ZETT (Zentrale Einrichtung für Tierforschung und wissenschaftliche Tierschutzaufgaben) under internal reference number O1016/25.

### Human pancreatic islets

Human islets were obtained from the NIDDK-funded Integrated Islet Distribution Program (IIDP; RRID:SCR_014387) at City of Hope, Duarte, CA, USA (NIH Grant U24DK098085). The use of human islets was approved by the Ethics Committee of the Medical Faculty, Heinrich Heine University Düsseldorf (study no. 3921). Islets from two non-diabetic donors were analyzed: a 66-year-old male (BMI 26.8 kg/m²; RRID:SAMN62478449) and a 54-year-old female (BMI 34.7 kg/m²; RRID:SAMN63285573).

### Islet isolation

Pancreatic islets were isolated from male C57BL/6J (Janvier) mice, age 9–11 weeks, as previously described, with minor modifications^117^. Following euthanasia, the common bile duct was clamped, and the pancreas was perfused with 3 ml of ice-cold Liberase™ RI (Roche, Cat. 11815032001) in DMEM (1×) + GlutaMAX™-I (Gibco™, Cat. 21885-035) via the bile duct. Perfused pancreata were excised and digested in a 37 °C water bath for up to 16.5 min. Digestion was stopped with DMEM containing 15% (v/v) FBS. Pancreata were mechanically dissociated by shaking and centrifuged (170x g, 3min). Pellets were resuspended and filtered through a sieve. Islets were purified by density-gradient centrifugation with DMEM + FBS over Lymphoprep™ (Serumwerk, Cat. 07851; 200× g, 25 min, no brake) and recovered from the interphase. Islets were handpicked under a stereomicroscope across two sequential washes and cultured overnight at 37 °C / 5% CO₂ in islet medium (CMRL 1066 [Gibco™, Cat. 21530-027] supplemented with 15% FBS, 1% penicillin/streptomycin, sodium bicarbonate, D-glucose, and 50 µM 2-mercaptoethanol) prior to treatment.

### Mass Spectrometry

Pancreatic islets were treated with 10 µM DXO (Sigma-Aldrich, Cat. D127) or a DXO derivative (Lam39M^28^) for 15 minutes. They were subsequently washed with 1 mL DPBS and centrifuged (500x g, 3 min, 37°C). The islets were solubilized in 200µl denaturing buffer (4% SDS, 100mM Tris, 20mM DTT, CompleteTM (Roche) and phosphatase Inhibitor cocktail (Sigma)). Lysis was performed by 10 strokes through an insulin syringe (26-gauge needle), followed by sonication (two cycles, pulse 0.09sec/10sec; Sonoplus, Bandelin). The lysates were than clarified by ultracentrifugation (75,300 x g, 30min, 4°C).

Protein concentrations in the supernatants were determined by direct photometric measurement (NanoDrop, Thermo Fisher Scientific). Approximately 100µg of protein was digested overnight at 37°C using Trypsin/LysC mix (1:25 (w/w) enzyme to protein ratio; Promega) on Mini S-Trap columns (ProtiFi) according to manufacturer’s instructions. Peptides were subsequently purified by solid-phase extraction (25µl RPS-Cartridges; Agilent) on an AssayMap Bravo platform (Agilent) following the supplier’s protocol.

Based on colorimetric peptide quantification (Pierce™), 45µg total peptide was used for phosphopeptide enrichment with FeNTA cartridges (5µl FeNTA-Cartridges; Agilent) on the AssayMap Bravo platform following the supplier’s protocol. Lyophilized peptides were reconstituted in 1% (v/v) trifluoroacetic acid (TFA) and analyzed on an Orbitrap Exploris 480 mass spectrometer coupled to a Vanquish Neo UHPLC (Thermo Fisher Scientific) operating in data-dependent acquisition (DDA) mode. Peptides were first trapped and desalted on an Acclaim PepMap C18 pre-column (75 µm inner diameter, 2 cm length; Thermo Fisher Scientific) and then separated on an Aurora C18 analytical column (25 cm × 75 µm, 1.6 µm particle size; IonOpticks, AUR2-25075C18A) at a flow rate of 300 nL/min. Chromatographic separation was performed at 40°C using a 2-hour, three-step linear gradient with mobile phase A (0.1% formic acid in water) and mobile phase B (80% acetonitrile, 0.1% formic acid in water), as follows:

0.5–19% buffer B over 72 minutes,

19–29% buffer B over 28 minutes,

29–41% buffer B over 20 minutes,

followed by a 1-minute ramp to 95% B.

Full MS scans were acquired at a resolution of 120,000 (at m/z 200) over an m/z range of 350–1,200, with a normalized AGC target of 300% and a 2-second cycle time. Precursors with charge states between 2+ and 6+ were selected for fragmentation, with a dynamic exclusion of 30 s.

MS/MS spectra were acquired at a resolution of 30,000 using higher-energy collisional dissociation (HCD) at a normalized collision energy of 30%. Fragment ions were isolated with a 1.6 m/z window, and AGC target values and injection times were automatically optimized (maximum injection time of 120 ms).

### Mass Spectrometry Data Analysis

Raw data were processed using Proteome Discoverer™ 3.1 (Thermo Fisher Scientific). Spectral recalibration was performed using the Spectrum RC node against the reviewed SwissProt Mus musculus (release v2023-11-08).

Label-free quantification was carried out using the Minora Feature Detector node with default settings, including a minimum trace length of 5, a maximum delta retention time of 0.2 min for isotope pattern multiplets, and identification restricted to high-confidence peptide-spectrum matches (PSMs).

Protein identification was performed using the Chimerys search engine (prediction model inferys_3.0.) against the reviewed SwissProt Mus musculus database (release v2024-03-27, including isoforms) supplemented with an in-house contaminant database. Trypsin was specified as the proteolytic enzyme, allowing up to two missed cleavages. Carbamidomethylation of cysteine was set as a static modification, while methionine oxidation and protein phosphorylation (Ser, Thr, Tyr) was set as a dynamic modification.

Label-free quantification was based on precursor ion intensities detected in at least 20% of the replicates. Normalization was performed on modified peptides (phosphorylated species) and peptide/protein abundance ratios were calculated using pairwise ratio-based comparisons with background-based t-tests.

Downstream analyses were done using in R (4.4.2) using RStudio (2026.01.0+392). The phosphopeptides with a peptide spectrum matching (PSM) count in the bottom 10nth percentile were removed to ensure detection quality. Subsequently, kinase perturbation analysis^39^ was performed with the perturbPlot2d function of the R package directPA within the PhosR workflow, using phosphosite-level. GO-term enrichment analysis was performed with the clusterProfiler^118^, enrichplot^119^, org.Mm.eg.db^120^, and AnnotationDbi^121^ packages. StringDB^122^ analysis was performed with the RStudio package STRINGdb with an interaction score cutoff of 750. Subsequently, refined by clustering networks utilizing Markov clustering (MCL), using an inflation parameter of 1.5. This allowed for in-depth analysis of the clusters, by the various enrichment tools STRINGdb offers (GO-term enrichment, Reactome pathways, KEGG pathways, WikiPathways). Function was inferred by these results, and the exported enrichment terms can be appreciated in supplemental table 4 and 5.

### Flow Cytometry

Per condition, 100 islets were treated with DXO (Sigma-Aldrich, Cat. D127), PD98059^63^ (Selleckchem, Cat. S1177), rapamycin^70^ (BLD-Pharm, Cat. BD33963), S961 acetate^71^ (MedChemExpress, Cat. HY-P2093B). Inhibitors were added 1h before DXO treatment was applied, STZ 24 h after DXO was added for an additional. Islets were collected, washed in PBS, and dispersed into a single-cell suspension with pre-warmed 0.05% trypsin-EDTA (Gibco, Cat. 25300054) by repeated pipetting (6–12 min). A heat-killed positive control was generated by incubation at 60 °C for 5 min. Cells were stained with Fixable Viability Stain 660 (BD Biosciences, Cat. 564405; 1:1000 in PBS) for 15 min at room temperature in the dark, with an unstained control included. Samples were acquired on a CytoFLEX S (Beckman Coulter); single, intact cells were gated by FSC-H/SSC-H and FSC-H/FSC-A, and viability was determined from the FVS660 (APC) channel, acquiring up to 10,000 events per sample. Data were analyzed in FlowJo^TM^ 10 (BD Biosciences).

### siRNA mediated knockdown in pseudo-islets

Pseudo-islets were generated from dissociated primary mouse or human islets to enable siRNA delivery, adapted from previous protocols^29,34,123^. Islets were dispersed into single cells with 0.05% trypsin-EDTA (Gibco, Cat. 25300-096), and cell number was determined utilizing a Neubauer chamber. Cells were transfected with 100 nM non-targeting control siRNA (si-*Ctrl*) or siRNA targeting a gene of interest (si-*Ror1*: ThermoFisher, s77260; si-*Ror2*: ThermoFisher, s77263; si-*ROR1*: ThermoFisher, s9755; si-*ROR2*: ThermoFisher, s9758; ON.TARGETplus non-targeting pool (si-*Ctrl* / si-*CTRL*): Horizon Discovery, D-001810-10-50) using Lipofectamine™ RNAiMAX (Thermo Fisher, Cat. 13778150) by transfection according to the manufacturer’s instructions. Transfected cells were seeded into conical 96-well plates (Sarstedt, Cat. 82.1583001) with 1500 cells per pseudo-islet and incubated for 24h at 37 °C / 5% CO₂ to allow aggregation and target knockdown. Where indicated, pseudo-islets were pre-treated with 10 µM DXO (Sigma-Aldrich, Cat. D127) for 24 h, followed by co-treatment with 1.5 mM or 15 mM streptozotocin (STZ; Sigma-Aldrich, Cat. S0130), mouse and human pseudo-islets respectively, for an additional 24 h. Knockdown efficiency was confirmed by qPCR and immunoblotting.

### Live cell imaging

Live/dead status was assessed in pseudo-islets by fluorescence microscopy. Pseudo-islets were stained in a dye solution containing Calcein-AM (2 µM), Hoechst 33342 (1:1000), and ethidium homodimer-1 (EthD; 4 µM) in KRH buffer supplemented with glucose (10 mM) and BSA and incubated at 37 °C for 60 min. Stained islets were transferred to a glass-bottom dish. Maximum intensity projections of z-stacks were acquired to obtain single islet images using an Apotome system, coupled to an Axio Observer Z1 microscope (Carl Zeiss Microscopy GmbH). Imaging was performed with a Plan-Apochromat 10×/0.45 objective. The area of dead cells compared to islet area was quantified in FIJI^124^ 2.16, using a previously established method^28^. Experiments with low knockdown efficiency (< 40% reduction either with or without DXO relative to si-*Ctrl*), or where STZ treatment was too harsh and did not allow proper image quantification were not included.

### Immunoblotting

Islets were collected in PBS, agglomerated (500x g, 3 min, 4 °C), and then lysed in RIPA buffer by mechanical disruption (10 min, 4 °C), and debris was pelleted by centrifugation (10,000× g, 10 min, 4 °C). Lysates were diluted with 4× Laemmli sample buffer (Bio-Rad, Cat. 1610747) and the proteins denatured at 95 °C for 10 min. Proteins were separated on 4–15% gradient SDS-PAGE gels (10 – 15 well gels, Bio-Rad), using 1x Tris/Glycine/SDS buffer (Bio-Rad, Cat. 1610772). Alongside a Precision Plus Protein™ All Blue Standard (Bio-Rad, Cat. 1610373) was used. Following electrophoresis, the proteins were transferred to membranes using the Trans-Blot Turbo system (Bio-Rad, 1704156). Membranes were blocked for 1 h at room temperature in 5% BSA or 5% milk in PBST and incubated overnight at 4 °C with primary antibodies (See Supplemental Table 9). After washing in PBST, membranes were incubated for 1 h at room temperature with HRP-conjugated anti-rabbit or anti-mouse IgG secondary antibody (See Supplemental Table 9; 1:1000–1:3000), developed with Clarity™ Western ECL substrate (Bio-Rad, Cat. 1705061) or SuperSignal^TM^ West Atto (ThermoFisher, Cat. A38555) and imaged on a ChemiDoc XRS (Bio-Rad). Antibodies are listed in supplemental table 9.

### Reverse transcription and quantitative PCR

Total RNA was isolated from islets using the RNeasy Mini Kit (Qiagen, 74104) with on-column DNase I digestion, per the manufacturer’s instructions. Islets were lysed in RLT buffer with mechanical disruption (10 min); RNA was eluted in RNase-free water. RNA was reverse-transcribed using the High-Capacity cDNA Reverse Transcription Kit (Thermo Fisher, Cat. 4368813) according to the manufacturer’s instructions. cDNA was diluted 1:3 in water for quantitative PCR (qPCR). qPCR was performed using Luna® Universal qPCR Master Mix (New England Biolabs, M3003). QuantStudioTM 1 System (Applied Biosystems) was used for experimental runs and analyzed in the QuantStudioTM Design and Analysis Desktop Software (Applied Biosystems). Primer sequences are presented in supplemental table 9.

### PamGene Analysis

Tyrosine kinase activity was profiled using the PamChip® PTK platform (PamGene International). Samples were treated with 10 µM DXO (Sigma-Aldrich, Cat. D127), DXO derivative (Kom56^30^), or (+)-Dizocilpine (MK801^76^) Maleate (Selleckchem, Cat. S2876). For kinase activity profiling, samples were pelleted (500× g, 8 min, 4 °C), washed in ice-cold PBS (1000× g, 5 min, 4 °C), and lysed on ice in in M-PER lysis buffer supplemented with protease and phosphatase inhibitor cocktails (1× final), with intermittent pipetting over 30 min, and cleared by centrifugation (10,000× g, 15 min, 4 °C). Lysates were cleared by centrifugation (10,000 × g, 15 min, 4 °C), snap-frozen, and stored at −80 °C; protein concentration was determined using the Pierce™ Bradford Plus Protein Assay Kit (Thermo Fisher, Cat. 23225) and 5 µg of total protein per array was used. Assays were run on a PamStation® 12 under Evolve software control. PTK PamChip® arrays were blocked with 2% BSA prior to sample application. A reaction mix containing PK buffer, BSA, DTT, ATP, PTK additive, and FITC-conjugated anti-phosphotyrosine detection antibody (PY20) (all supplied in the PamGene assay Kit) was combined with the diluted lysate, and 40 µl of the complete mix was applied per array. Peptide phosphorylation was measured in real time by imaging the fluorescently labelled antibody bound to the 196 immobilized tyrosine-kinase substrate peptides on each array according to the manufacturer.

### Quantification and statistical analysis

Protein quantification and statistical analysis were performed in Proteome Discoverer version 3.0 (Thermo Fisher Scientific). Protein-level fold changes between control and DXO and DXO derivative treated islets were calculated were calculated using pairwise ratio-based comparisons. Statistical significance was assessed using background-based *t*-tests implemented in Proteome Discoverer. *P* values were adjusted for multiple testing using the Benjamini–Hochberg method.

Statistical analyses were otherwise performed utilizing GraphPad Prism 11.1.0, as indicated in the figure legends. Data are reported as mean ± SEM unless stated otherwise. Microscopy-based assessments of cell viability were conducted with the investigator blinded to the experimental conditions. Number of biological replicates and independent experiments are described in the figure legends.

---

**Supplemental Figure 1:**
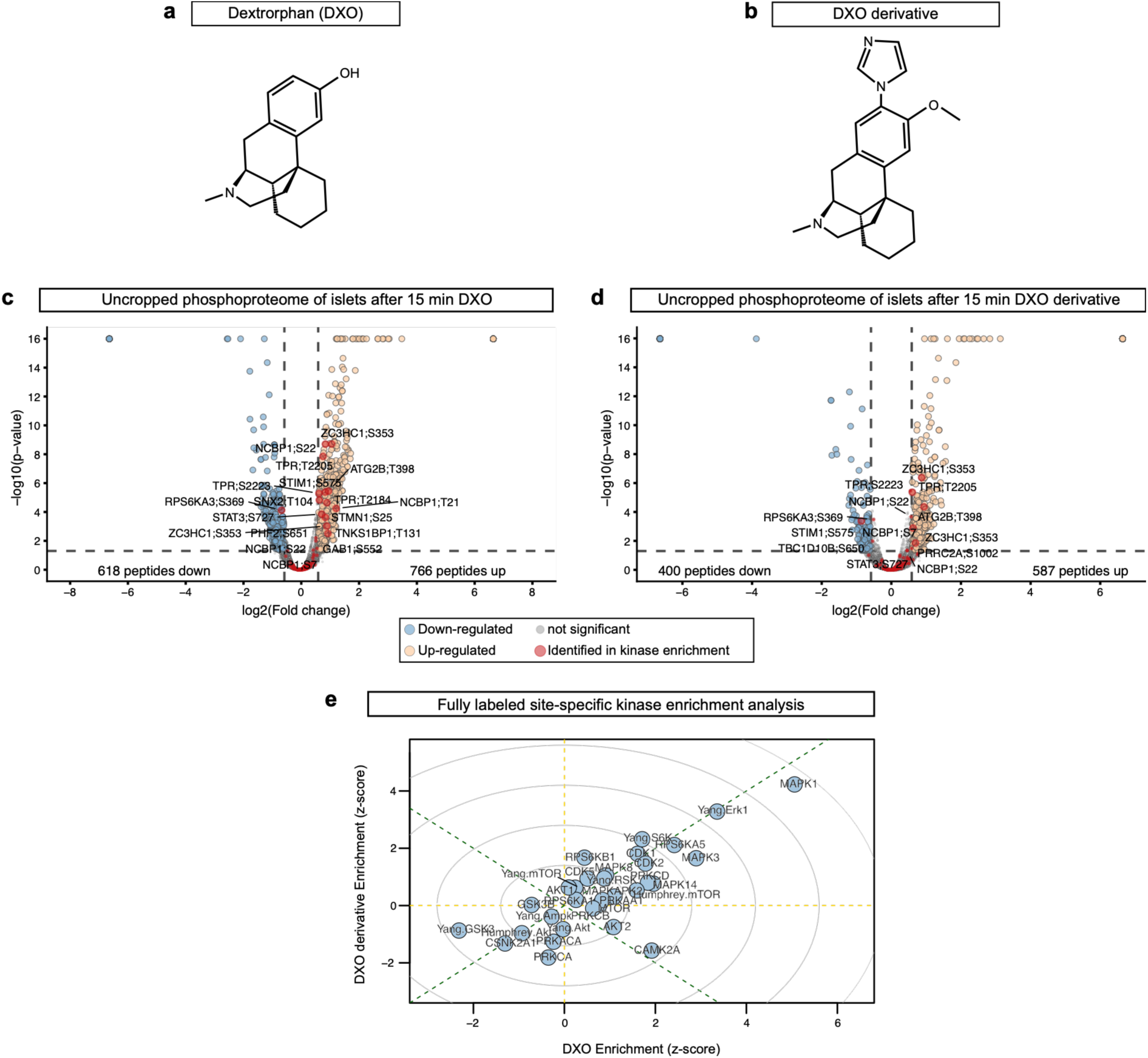
Full phosphoproteomics volcano plots and kinase enrichment analysis. (a, **b)** Chemical structure of dextrorphan (a) and BBB-impermeable DXO derivative^28^ (b). **(c, d)** Uncropped volcano plots of 15 min DXO (c) or DXO derivative (d) treated pancreatic islets compared to vehicle control. Orange dots mark significantly upregulated, blue dots significantly downregulated, and grey dots unchanged phosphopeptides (|log2(FC)| > 0.585;-log10(p-value) > 1.301, as indicated by dashed lines). Red dots highlight phosphopeptides associated with signaling pathways identified in downstream kinase enrichment analyses (e) (n = 6 islet preparations). **(e)** Kinase enrichment analysis of Fig. 1d with all data points labeled. Different datasets that were used for enrichment analysis are labeled as such.

**Supplemental Figure 2:**
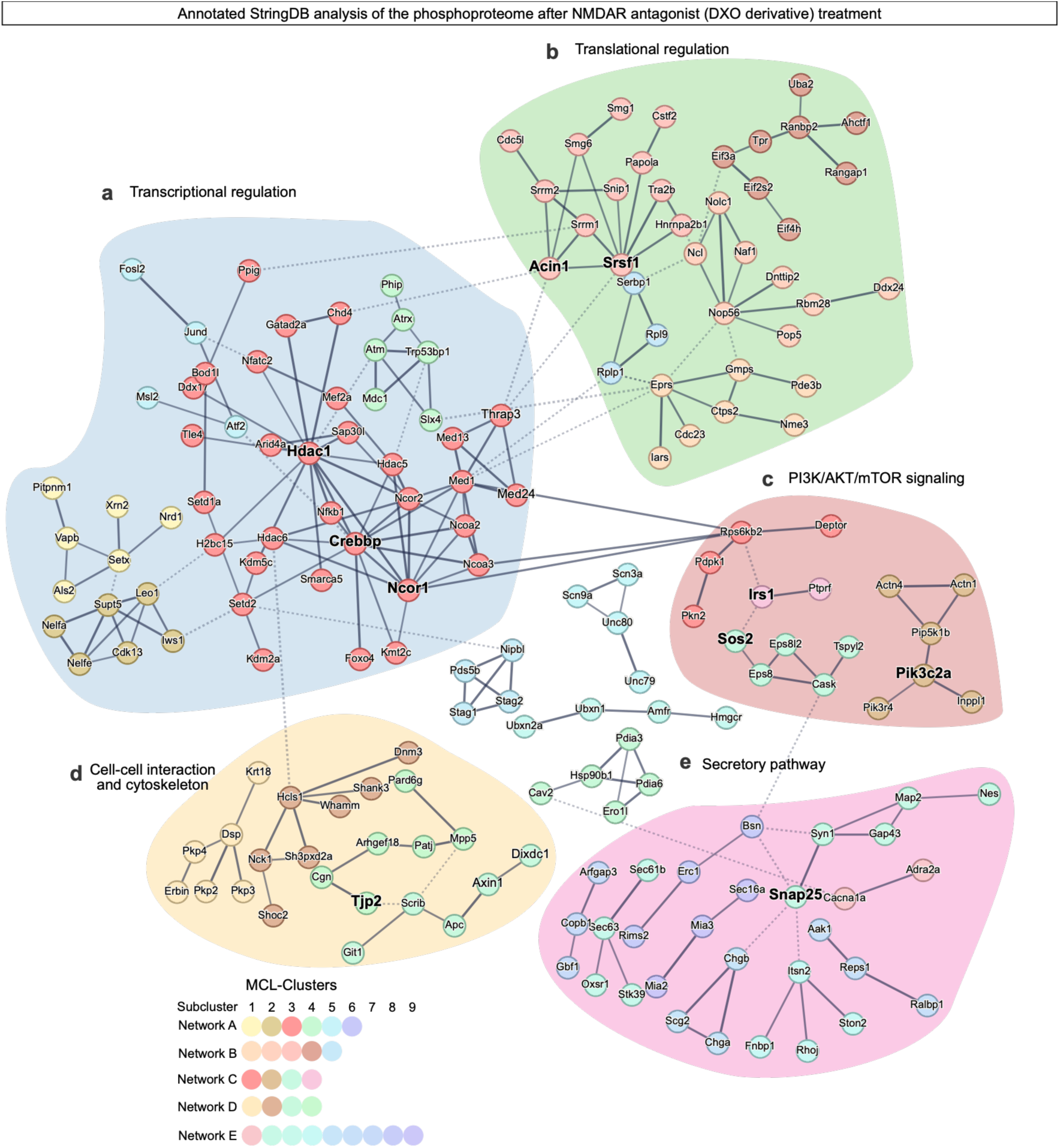
StringDB analyses of differentially regulated phosphopeptides after treating islets with BBB-impermeable DXO derivative. Annotated protein high-confidence interaction network of significantly altered phosphopeptides in pancreatic islets after 15 min treatment with NMDA receptor antagonist DXO derivative (StringDB package in RStudio; score threshold = 750; 222 phosphopeptides total). Disconnected nodes and small networks were omitted for clarity. Connected nodes were partitioned using the Markov clustering algorithm (MCL, inflation parameter = 1.5; each cluster was assigned a color). MCL-clusters were arranged according to the enriched GO-terms of the underlying clusters to gain insights into functional connections, which resulted in the colored fields and respective titles (Table S5): **(a)** Transcriptional regulation; **(b)** Translational regulation; **(c)** PI3K/AKT/mTOR signaling; **(d)** Cell-cell interaction and cytoskeleton; and **(e)** Secretory pathway.

**Supplemental Figure 3:**
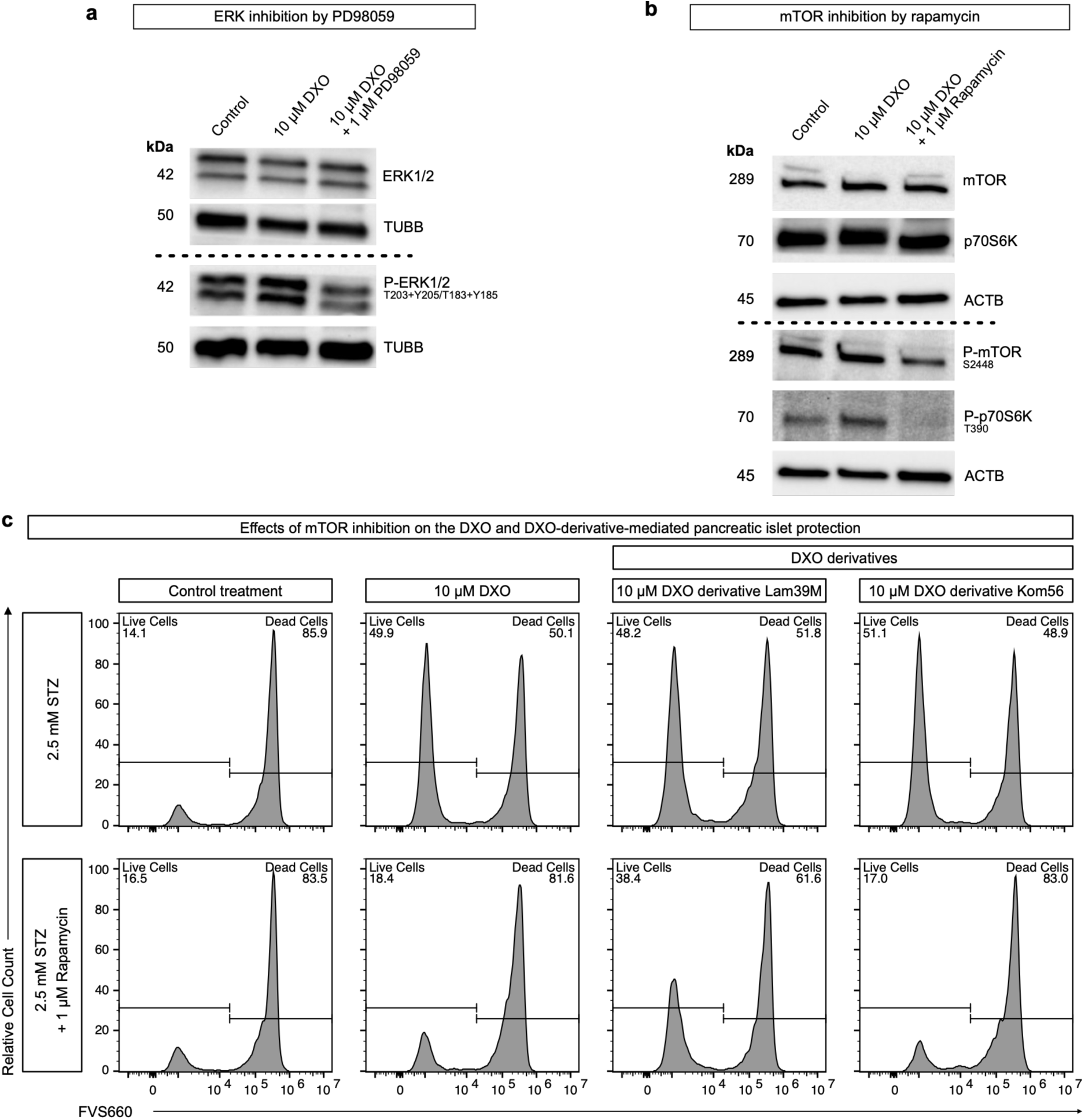
Effects of mTOR inhibitor rapamycin on the islet cell-protective effects of DXO and its derivatives. (a, **b)** Immunoblot with 10 µM DXO ± 1 µM PD98059 (a) or 1 µM rapamycin (b) to show effective inhibition ERK1/2 and mTOR, respectively. ERK1/2, P-ERK1/2 (T203+Y205/T183+Y185), and TUBB are shown in (a). mTOR, p70S6K, P-mTOR (S2448), P-p70S6K (T390), and ACTB are shown in (b). TUBB and ACTB were used as loading controls (a, b respectively). **(c)** Flow cytometry experiment of pancreatic islets treated with 10 µM DXO and DXO derivatives Lam39M^28^ or Kom56^30^ with or without 1 µM rapamycin (48 h) ± 2.5 mM STZ (24 h).

**Supplemental Figure 4:**
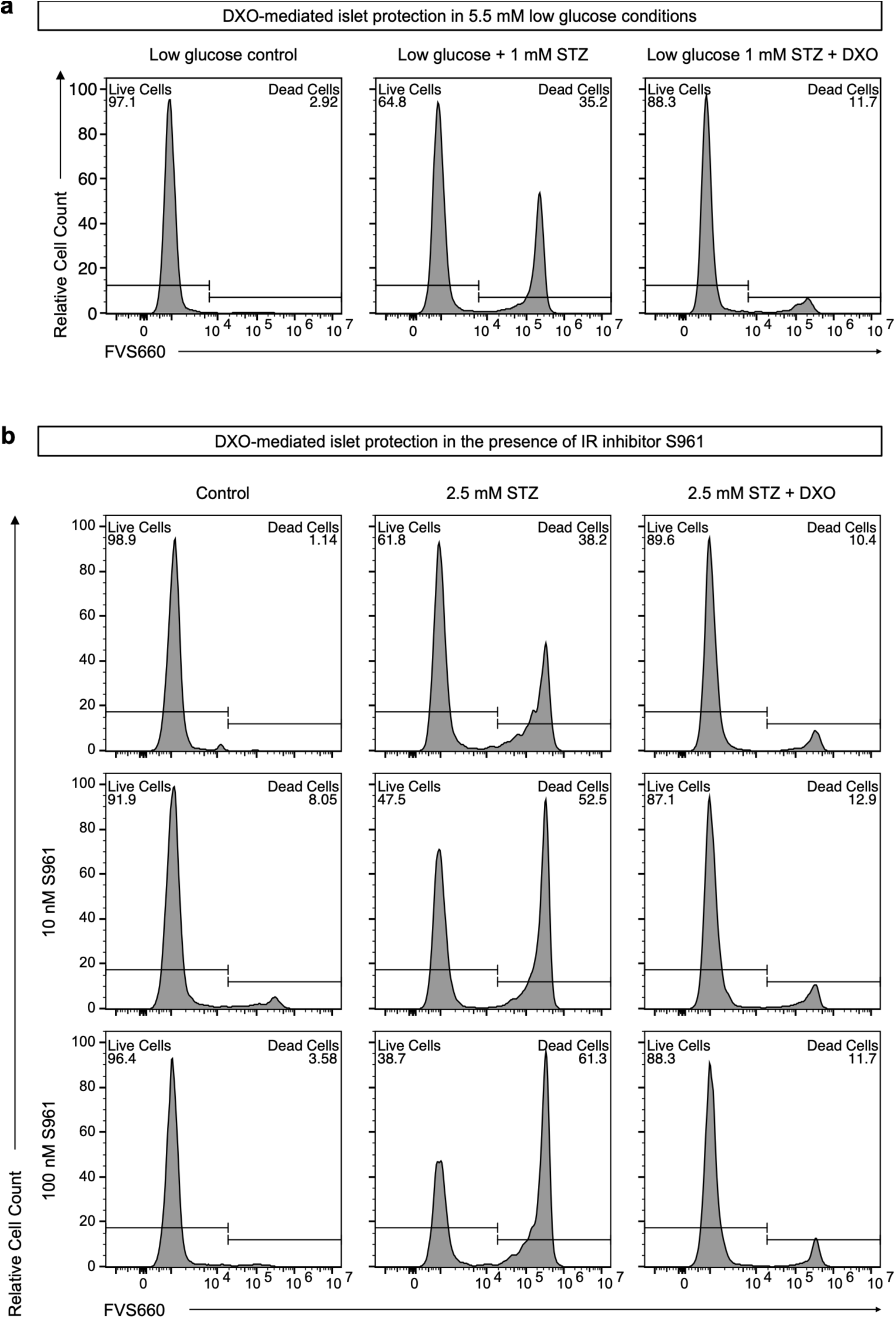
Islet cell-protective effects of DXO require neither glucose stimulation nor insulin receptor activation. **(a)** Flow cytometry experiment of pancreatic islets ± 1 mM STZ (24 h) with or without 10 µM DXO (48 h) in 5.5 mM glucose medium (low glucose). **(b)** Flow cytometry experiment of pancreatic islet cells treated with 10 µM DXO with or without 10 nM or 100 nM insulin receptor (IR) inhibitor S961^115^ (48 h) ± STZ (24 h).

**Supplemental Figure 5:**
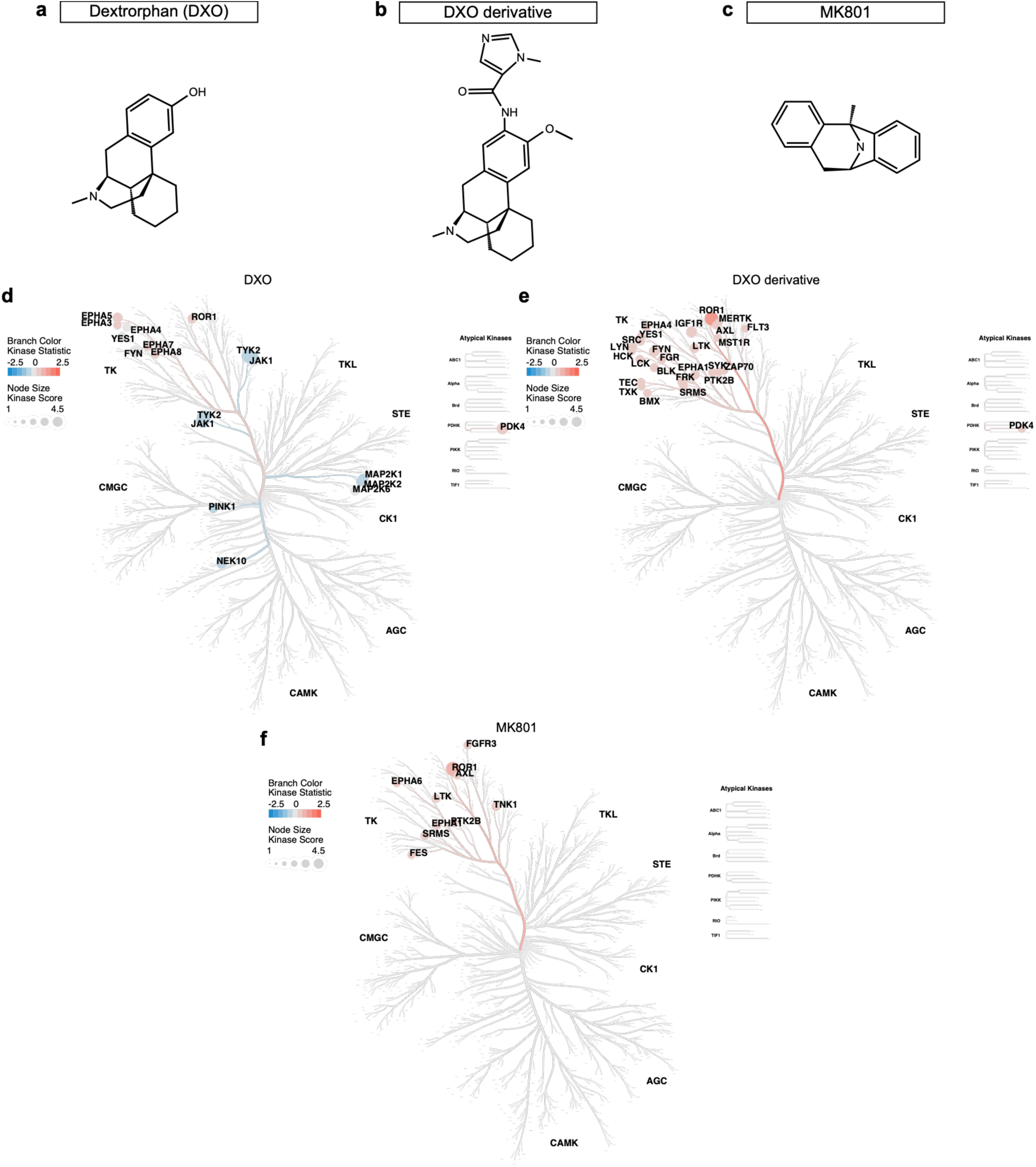
Uncropped phylogenetic CORAL kinome trees after treating islets with three different NMDAR-antagonists shown in. Fig. 4b**-d**. Kinome activity profile visualized on a CORAL kinome tree. Pancreatic islets were treated with 10 µM DXO **(a, d)**, 10 µM BBB-impermeable DXO derivative^30^ **(b, e)**, and 10 µM NMDA receptor-selective antagonist MK801^21,76^ **(c, f)** for 15 minutes and later analyzed using the PamGene assay kit. In the CORAL kinome trees, the branch color indicates the “kinase statistic” (up-or downregulation) and node size depicts the “kinase score” (a measure of significance). Here, the full phylogenetic tree is shown, while the main figure was reduced to the tyrosine kinase (TK) branch.

**Supplemental Figure 6:**
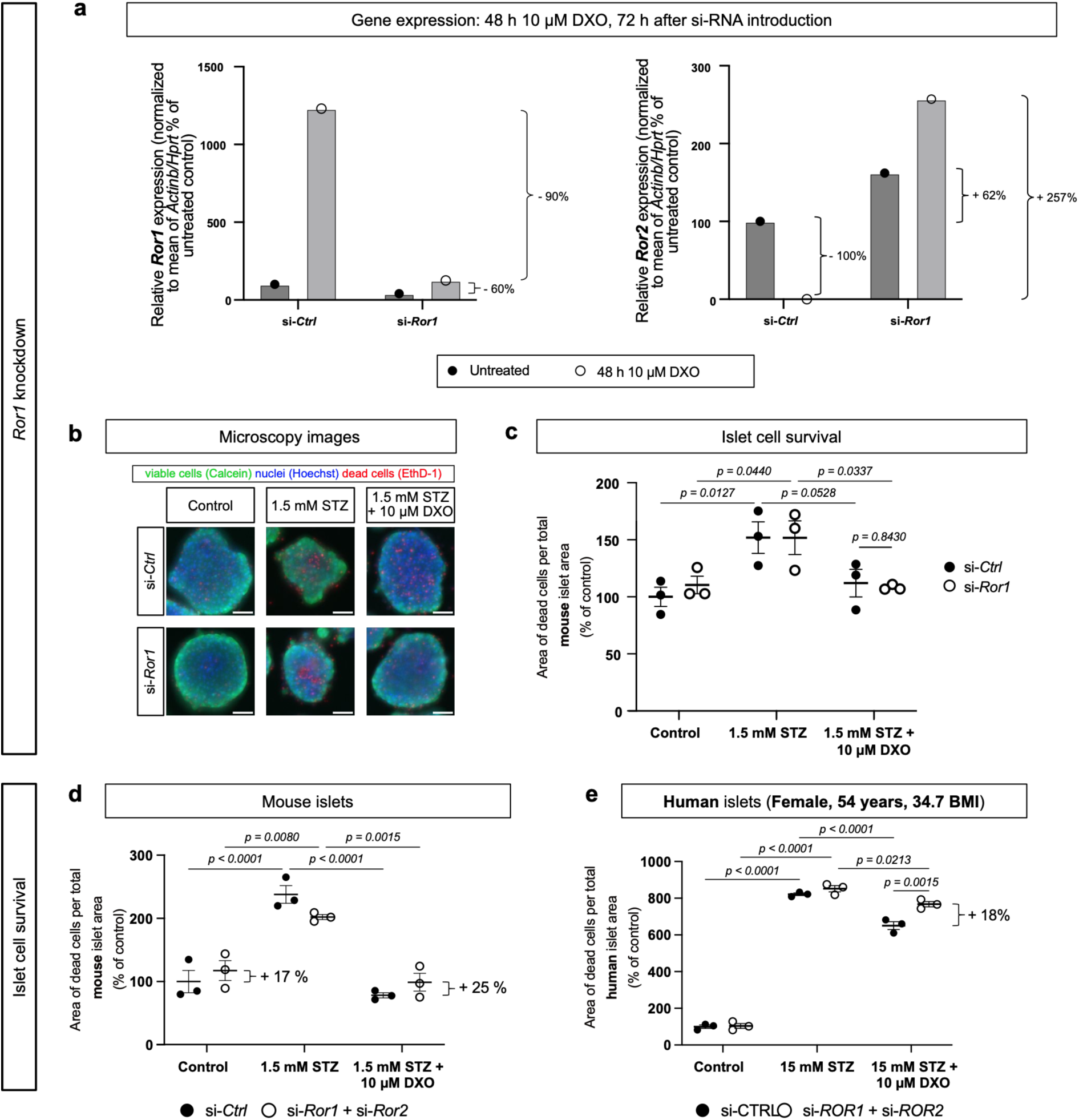
*Ror1* knockdown does not ameliorate DXO-mediated islet cell protection. **(a)** Relative mRNA expression of *Ror1* in si-*Ror1* versus si-*Ctrl* siRNA transfected mouse pseudo-islets treated with or without 10 µM DXO (48 h). Data are shown as individual values normalized to the housekeeping mRNA *Actb* and *Hprt* relative to si-*Ctrl* siRNA (n = 1). **(b)** Images of live-dead imaging of si-*Ctrl* vs si-*Ror1* siRNA transfected mouse pseudo-islets treated with or without 1.5 mM STZ (24 h) ± 10 μM DXO (48 h). Nuclei were stained with Hoechst (blue color), viable cells with calcein (green color), and dead cells with ethidium-homodimer-1 (Ethd-1; red color). **(c)** Quantification of the area of dead cells of the experiment shown in (b). **(d)** Replicate of the si-*Ror1* + si-*Ror2* knockdown in mouse pseudo-islets shown in Fig. 5d, e. **(e)** Replicate of the si-*ROR1* + si-*ROR2* knockdown in human pseudo-islets shown in Fig. 5g, h. A repeated measures two-way ANOVA with Sidak’s post hoc test was performed to test for statistical significance. Data points are presented as mean ± SEM (c-e).

**Appendix Figure 3.I:**
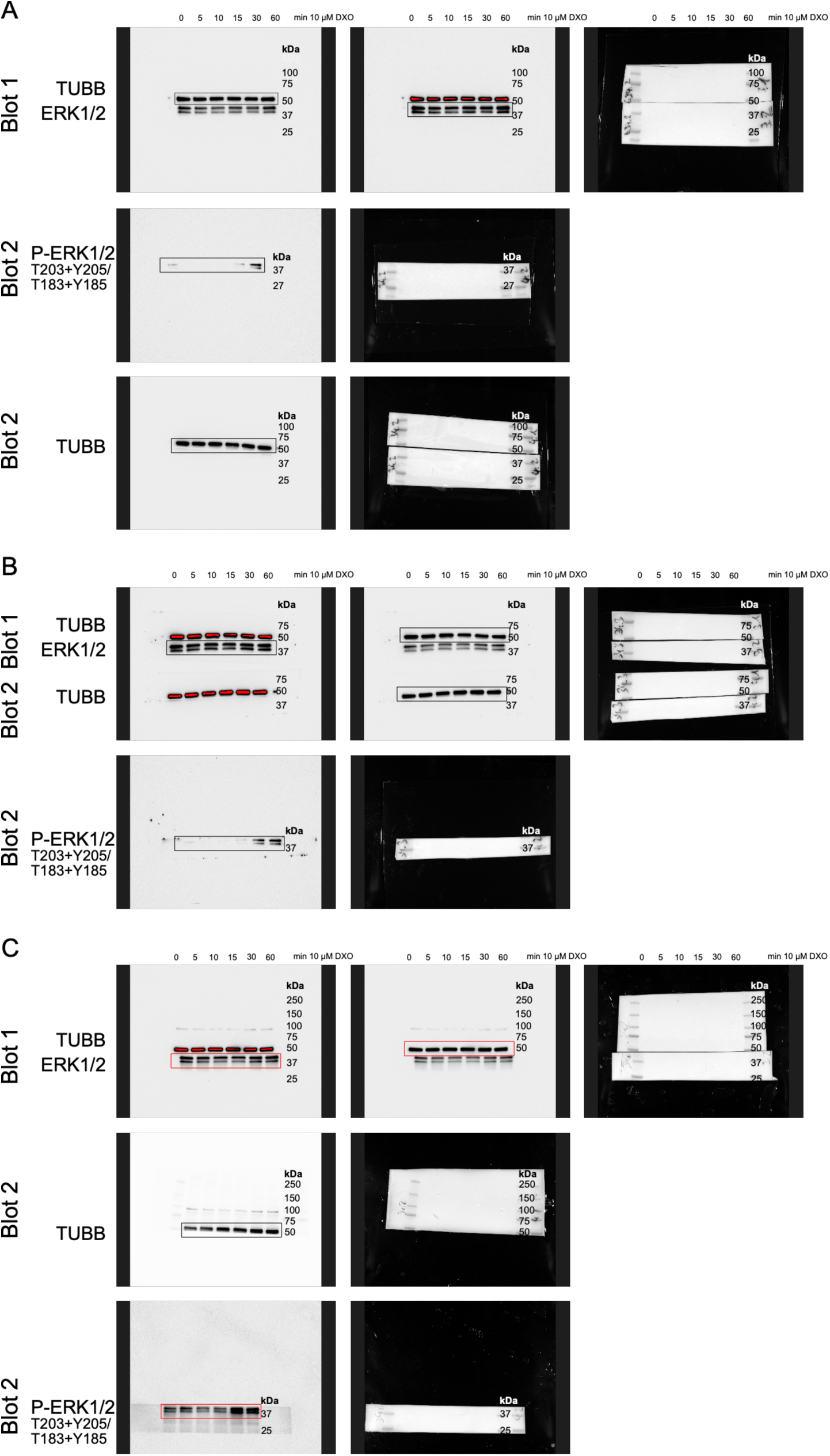
Uncropped immunoblot images related to. **Fig. 3a**. Time-course immunoblots of pancreatic islets treated with 10 µM DXO for 0, 5, 10, 15, 30, and 60 minutes. Total ERK1/2 and p-ERK1/2 (T203+Y205/T183+Y185) are shown, and TUBB was used as a loading control. The pan-protein blots and the phospho-protein blots are shown (distinguishable by Blot 1/2 label) as chemiluminescence images and the corresponding colorimetric images (with molecular weight markers). Three independent replicates were performed **(A-C)**. Black boxes indicate bands that were used for analysis, red boxes indicate the images shown in the main figure.

**Appendix Figure 3.II:**
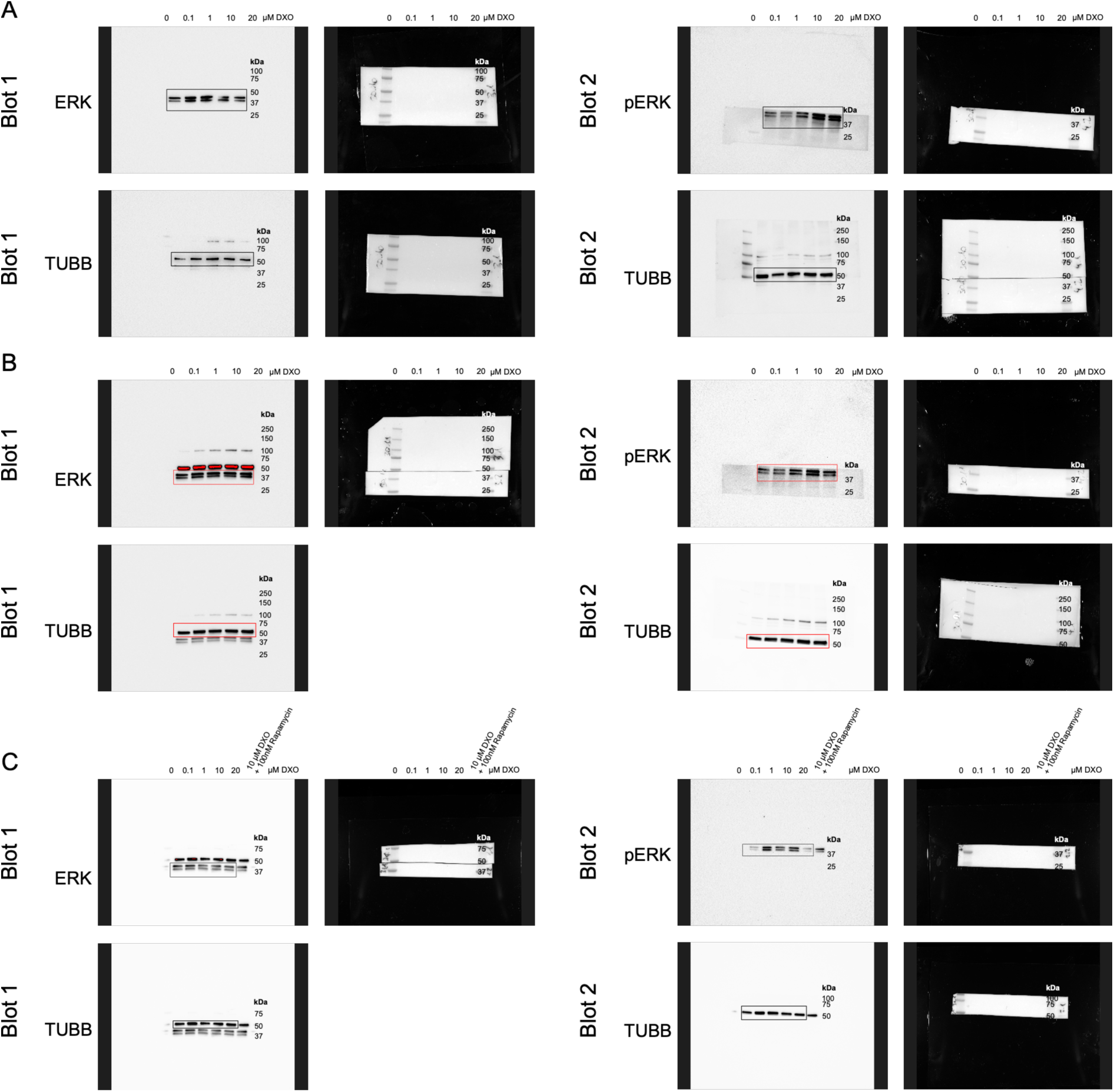
Uncropped immunoblot images related to. **Fig. 3b**. Dose-response immunoblots of pancreatic islets treated with different DXO concentrations for 24 h. Total ERK1/2 and p-ERK1/2 (T203+Y205/T183+Y185) are shown, and TUBB was used as a loading control. The pan-protein blot (Blot 1) and the phospho-protein blot (Blot 2) are shown as chemiluminescence images and the corresponding colorimetric images (with molecular weight markers). Three independent replicates were performed **(A-C)**. Black boxes indicate bands that were used for analysis, red boxes indicate the images shown in the main figure.

**Appendix Figure 3.III:**
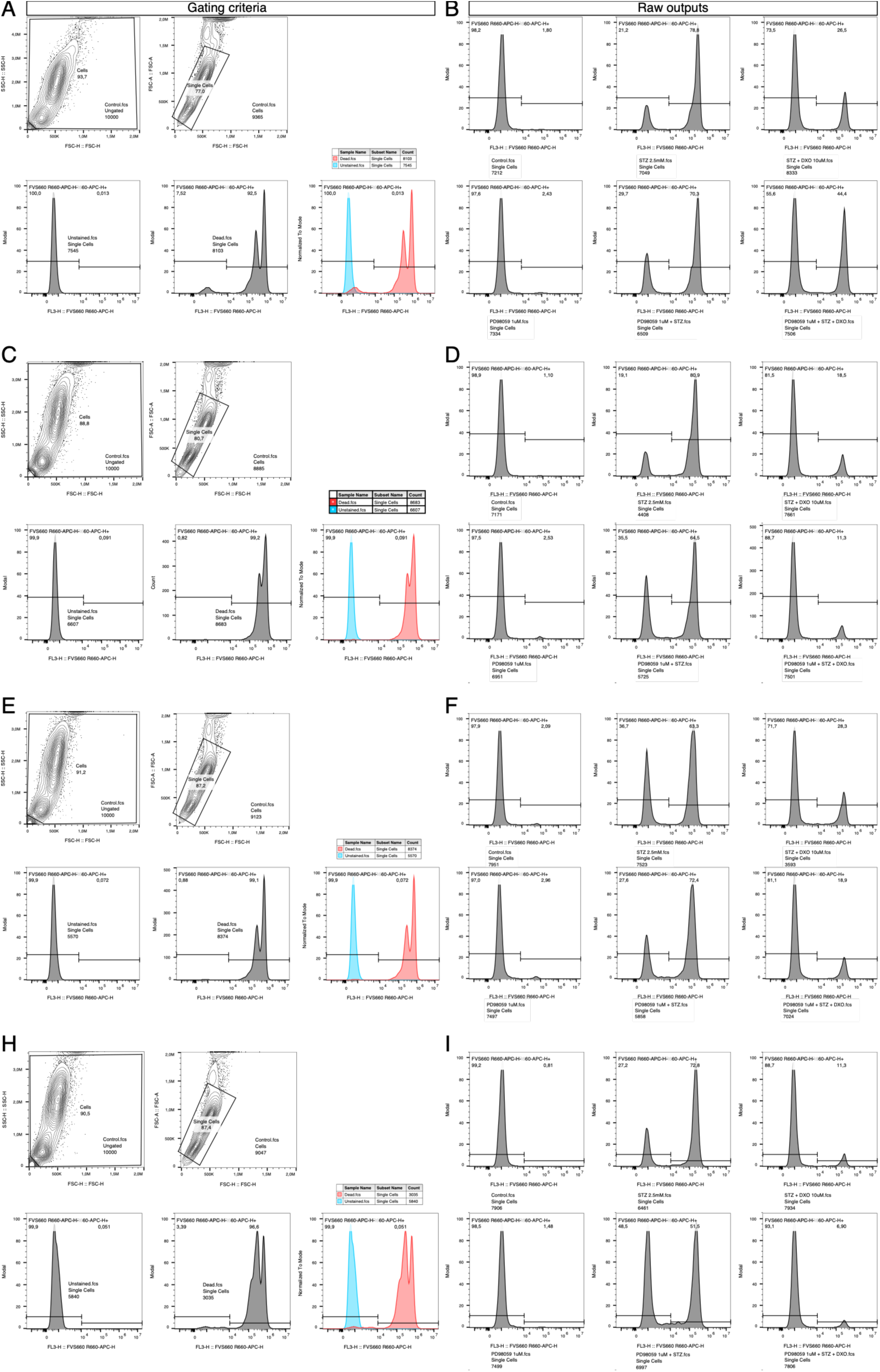
Gating criteria and raw output of the flow cytometry measurement of. **Fig. 3c-d. (A)** Gating criteria for flow cytometry analysis of the first replicate of the analysis in the main figure. Cell debris were excluded using FSC-H – SSC-H projection, resulting in “Cells”. “Single Cells” were gated from “Cells” using FSC-A – FSC-H. “Single Cells” were then used to distinguish dead from living cells in the APC (FVS660) channel. **(B)** Raw outputs of the first replicate of the flow cytometric measurements before rearranging plots for the main figure. **(C, D)** represents the second replicate. **(E, F)** represents the third (representative) replicate, which is shown in Fig. 3c. **(H, I)** represents the fourth replicate.

**Appendix Figure 3.IV:**
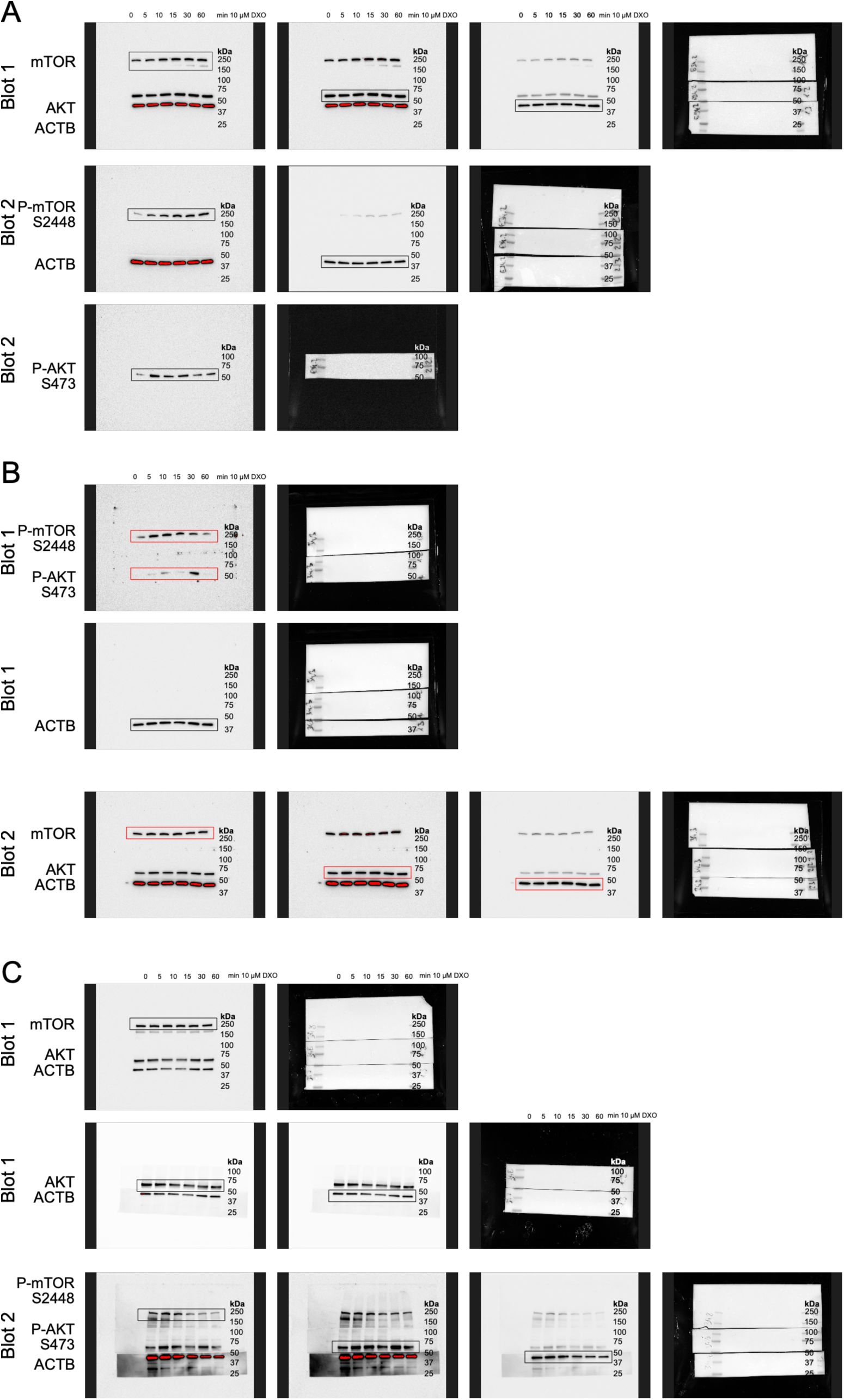
Uncropped immunoblot images related to. **Fig. 3e**. Time-course immunoblots of mouse islets treated with 10 µM DXO for 0, 5, 10, 15, 30, and 60 minutes. Total mTOR, p-mTOR (S2448), total AKT and p-AKT (S473) are shown, and ACTB was used as a loading control. The pan-protein blots and the phospho-protein blots are shown (distinguishable by Blot 1/2 label) as chemiluminescence images and the corresponding colorimetric images. Three independent replicates were performed **(A-C)**. Black boxes indicate bands that were used for analysis, red boxes indicate the images shown in the main figure.

**Appendix Figure 3.V:**
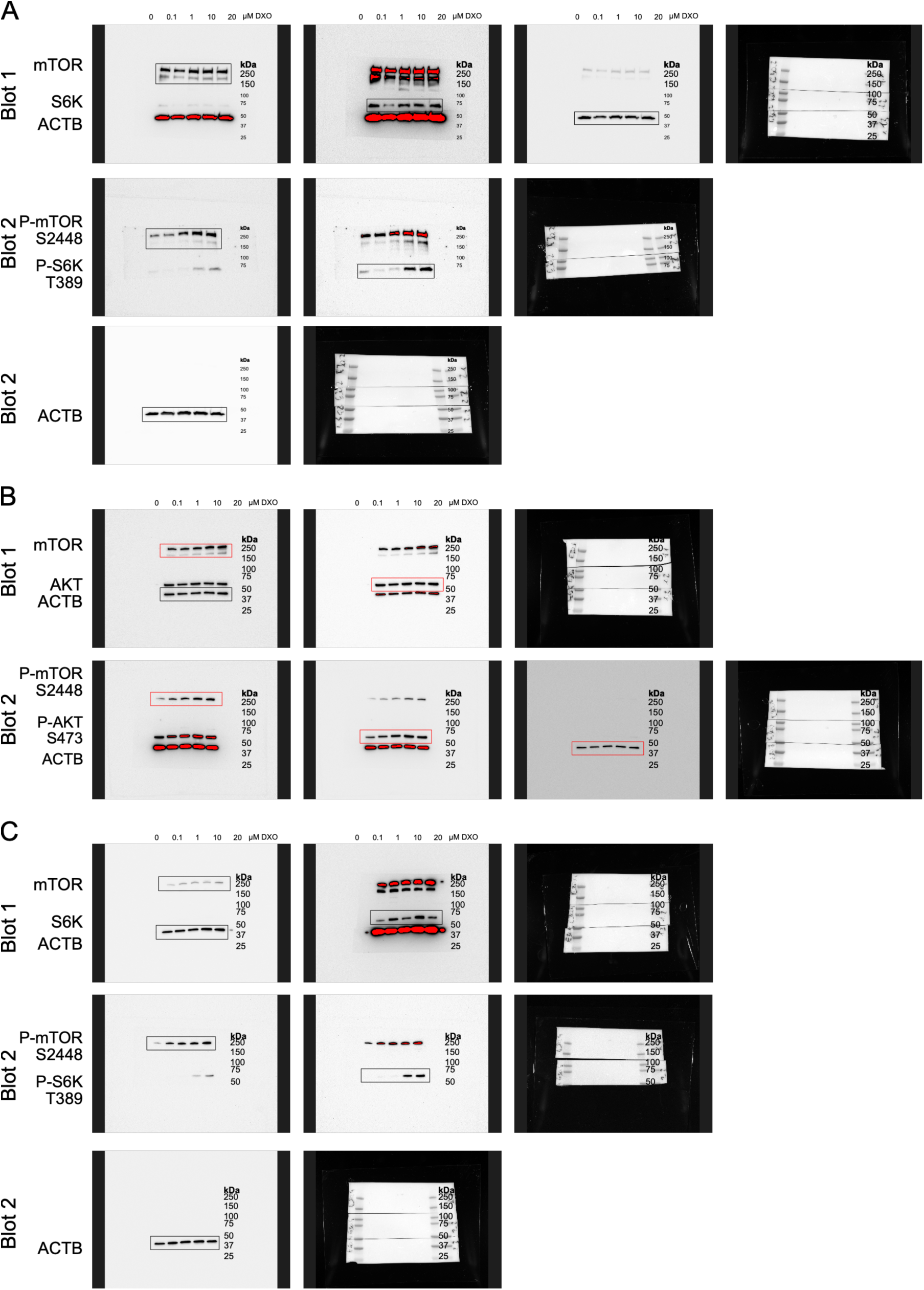
Uncropped immunoblot images related to. **Fig. 3f**. Dose-response immunoblots of pancreatic islets treated with 0, 0.1, 1, 10, 20 µM DXO for 24 h. Total mTOR, p-mTOR (S2448), total AKT and p-AKT (S473) are shown, and ACTB was used as a loading control. The pan-protein blots and the phospho-protein blots are shown (distinguishable by Blot 1/2 label) as chemiluminescence images and the corresponding colorimetric images (with molecular weight markers). Five independent replicates were performed **(A-C)**, continued in App. Fig. 3.VI. Black boxes indicate bands that were used for analysis, red boxes indicate the images shown in the main figure.

**Appendix Figure 3.VI:**
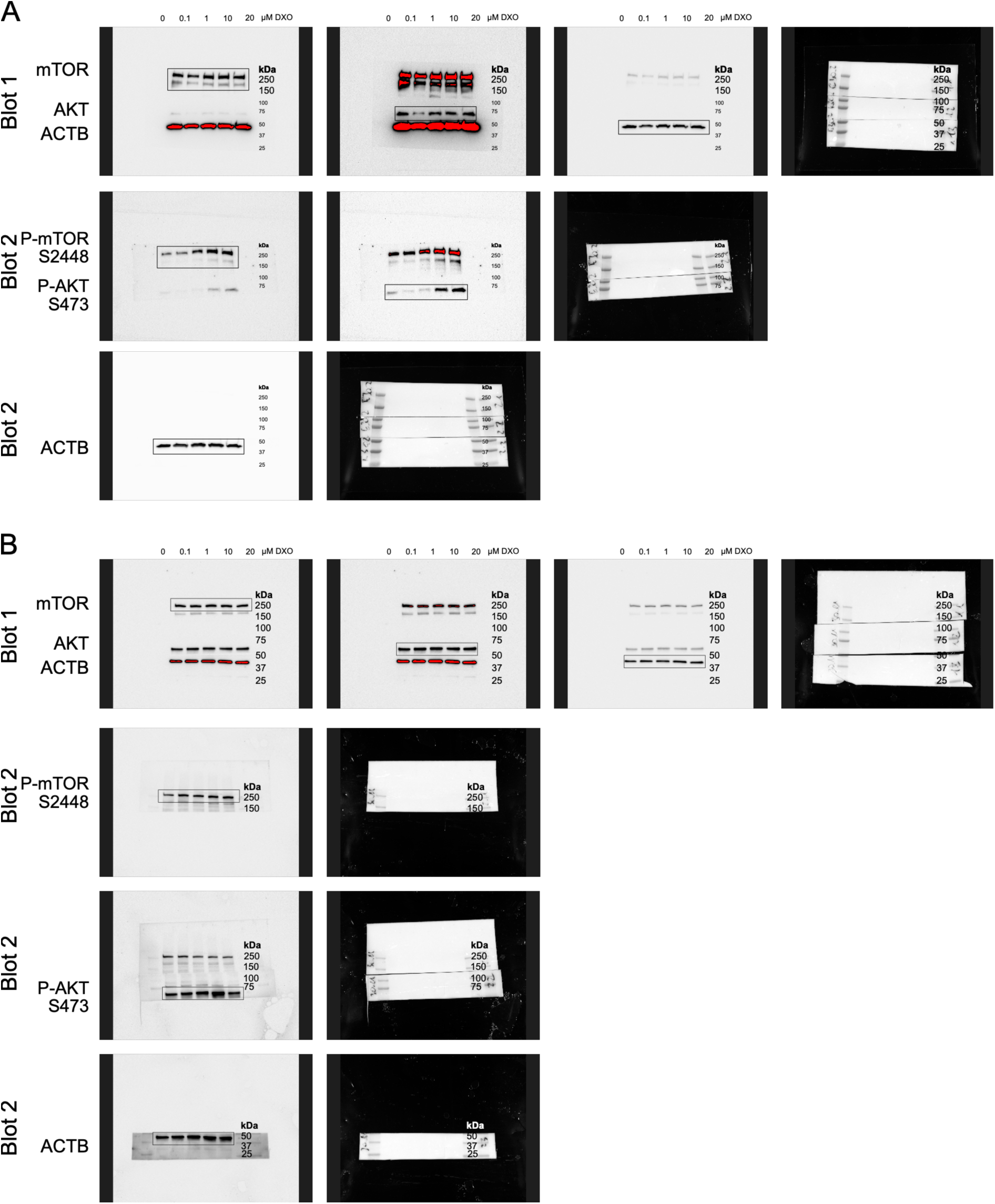
Uncropped immunoblot images related to. **Fig. 3f**. Dose-response immunoblots of pancreatic islets treated with 0, 0.1, 1, 10, 20 µM DXO for 24 h. Total mTOR, p-mTOR (S2448), total AKT and p-AKT (S473) are shown, and ACTB was used as a loading control. The pan-protein blots and the phospho-protein blots are shown (distinguishable by Blot 1/2 label) as chemiluminescence images and the corresponding colorimetric images. Continued from App. Fig. 3.V **(A-B)** are the final two replicates. Black boxes indicate bands that were used for analysis.

**Appendix Figure 3.VII:**
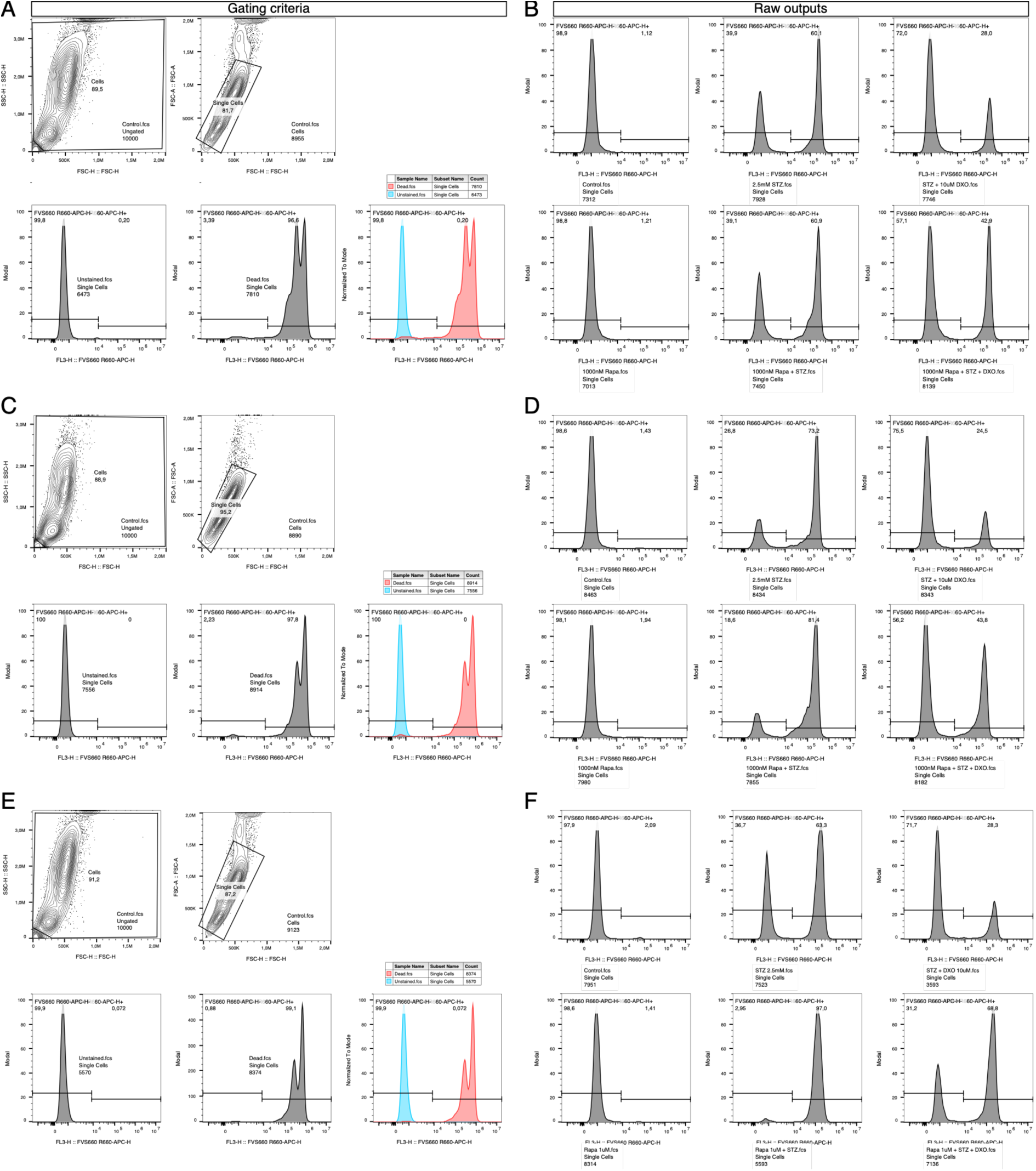
Gating criteria and raw output of the flow cytometry measurement of. **Fig. 3G-H. (A)** Gating criteria for flow cytometry analysis of the first replicate of the analysis in the main figure. Cell debris were excluded using FSC-H – SSC-H projection, resulting in “Cells”. “Single Cells” were gated from “Cells” using FSC-A – FSC-H. “Single Cells” were then used to distinguish dead from living cells in the APC (FVS660) channel. **(B)** Raw outputs of the first replicate of the flow cytometric measurement before rearranging plots for the main figure. **(A, B)** represents the representative experiment shown in Fig. 4C. **(C, D)** represents the second replicate. **(E, F)** represents the third replicate.

**Appendix Figure S3.I:**
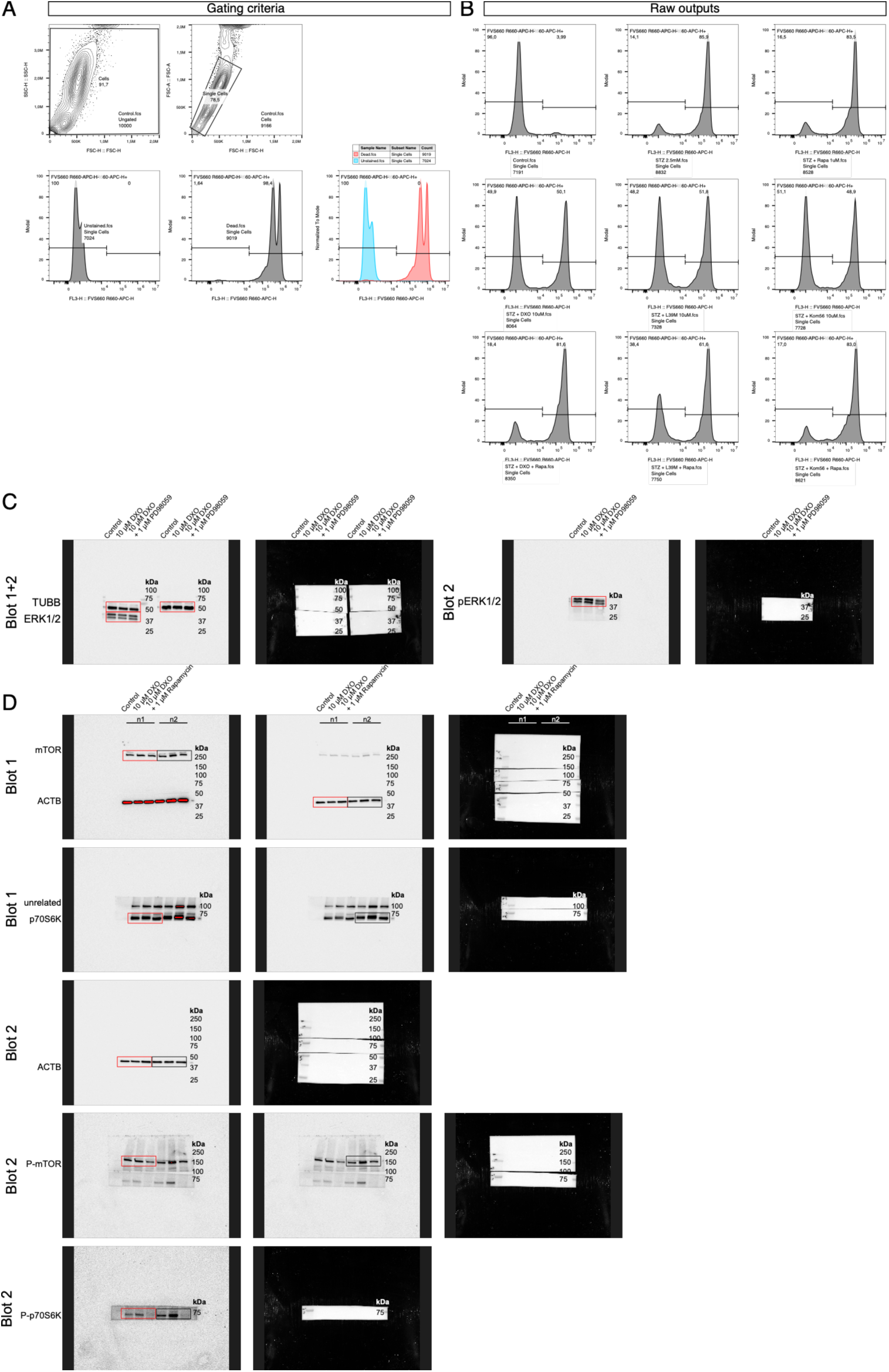
Flow cytometry and uncropped immunoblot images related to supplemental Fig. 3. **(A)** Gating criteria for flow cytometry analysis of the first replicate of the analysis in the main figure. Cell debris were excluded using FSC-H – SSC-H projection, resulting in “Cells”. “Single Cells” were gated from “Cells” using FSC-A – FSC-H. “Single Cells” were then used to distinguish dead from living cells in the APC (FVS660) channel. **(B)** Raw outputs of the flow cytometric measurement before rearranging plots for the main figure. **(C+D)** Immunoblots of pancreatic islets treated 1h of either 1µM MEK-inhibitor PD98059 (C) or 1 µM of mTOR inhibitor rapamycin for 1h before addition of 10 µM DXO for 24 h. (C) Total ERK1/2 and p-ERK1/2 (T203+Y205/T183+Y185) are shown, and TUBB was used as a loading control. (D) Total mTOR, p-mTOR (S2448), total p70S6K and p-p70S6K (T390) are shown, and ACTB was used as a loading control. The pan-protein blots and the phospho-protein blots are shown (distinguishable by Blot 1/2 label) as chemiluminescence images and the corresponding colorimetric images. Black boxes indicate bands that were used for analysis, red boxes indicate the images shown in the main figure.

**Appendix Figure S4.I:**
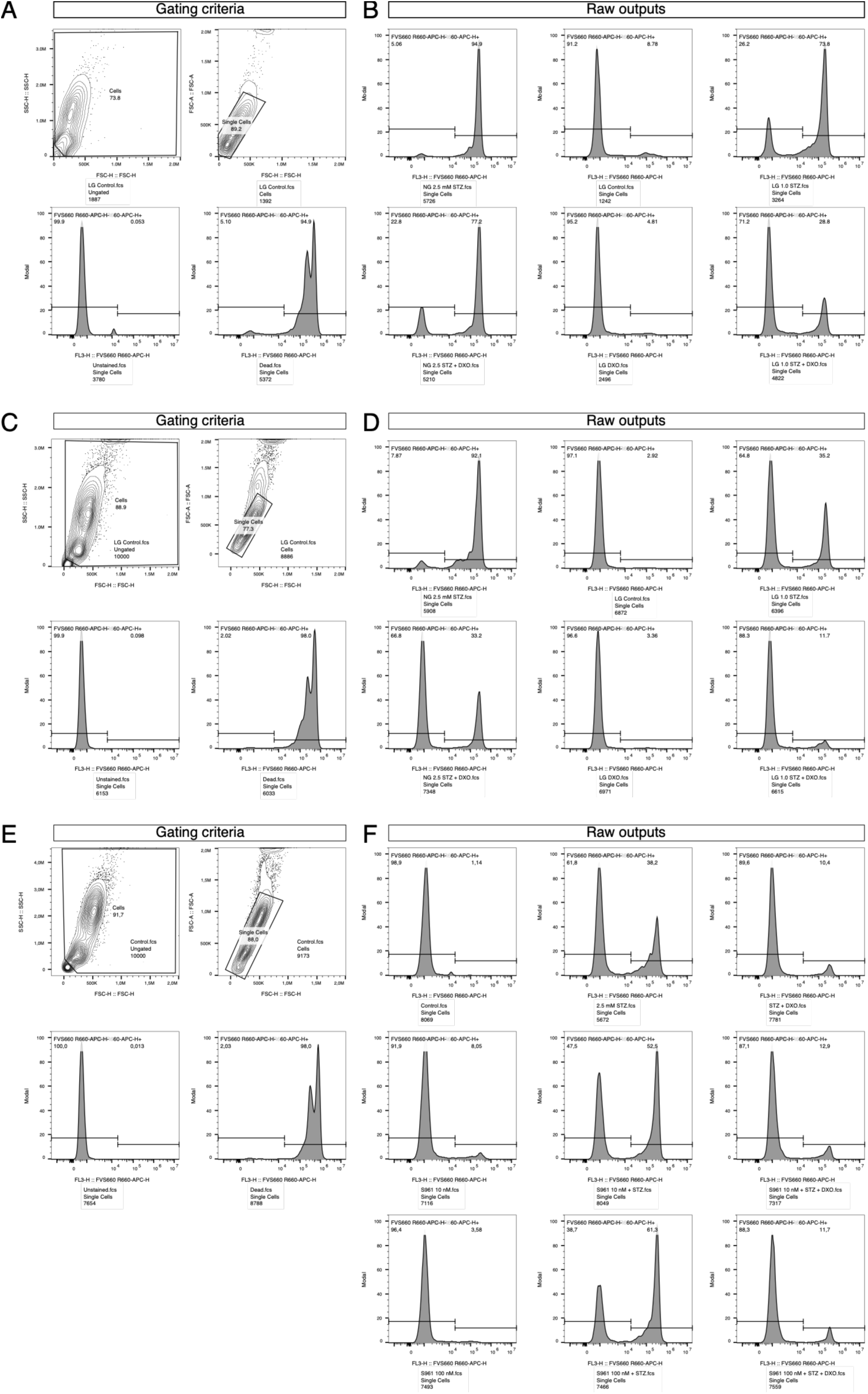
Flow cytometry related to supplemental Fig. 4. (A, C, **D)** Gating criteria for flow cytometry analysis of the first replicate of the analysis in the main figure. Cell debris were excluded using FSC-H – SSC-H projection, resulting in “Cells”. “Single Cells” were gated from “Cells” using FSC-A – FSC-H. “Single Cells” were then used to distinguish dead from living cells in the APC (FVS660) channel. **(B, D, F)** Raw outputs of the flow cytometric measurement before rearranging plots for the main figure. (A-D) are related to supplemental Fig. 4a and represent two independent replicates, while (C, D) represent the experiment shown in the main figure. (E+F) are related to supplemental Fig. 4b.

**Supplemental Table 1: Raw data of the phosphoproteome measurements**. Here, the raw output after mass spectrometry analysis is shown. This was used as the input data for downstream statistical analyses shown in Fig. 1 and Fig 2. n = 6 independent experiments. Conditions: Control, 10 µM DXO, 10 µM BBB-impermeable DXO-derivative for 15 min.

**Supplemental Table 2**: Full GO term enrichment analysis results of DXO-induced phosphoproteome changes as shown in Fig. 1e.

**Supplemental Table 3**: Full GO term enrichment analysis results of DXO derivative-induced phosphoproteome changes as shown in Fig. 1e.

**Supplemental Table 4**: The GO term enrichment results of the DXO-induced phosphoproteome derived STRING clusters in Fig. 2. The top 5 (or less) of enrichment results are listed here. Titles of subnetworks in Fig. 2 were inferred from these analyses.

**Supplemental Table 5**: The GO term enrichment results of the DXO derivative-induced phosphoproteome derived STRING clusters in supplemental figure 2. The top 5 (or less) of enrichment results are listed here. Titles of subnetworks in supplemental figure 2 were inferred from these analyses.

**Supplemental Table 6**: PamGene Analysis output data of DXO-treated islets. The list contains comparisons between the different treatment groups. Statistical values were assigned to each kinase, as they were only partly displayed (in case of change in activity) in Fig. 4b.

**Supplemental Table 7**: PamGene Analysis output data of DXO derivative-treated islets. The list contains comparisons between the different treatment groups. Statistical values were assigned to each kinase, as they were only partly displayed (in case of change in activity) in Fig. 4c.

**Supplemental Table 8**: PamGene Analysis output data of MK801-treated islets. The list contains comparisons between the different treatment groups. Statistical values were assigned to each kinase, as they were only partly displayed (in case of change in activity) in Fig. 4d.

**Supplemental Table 9:** Primary and Secondary Antibodies and primers used in this study.

## Literature

1 Abel, E. D., et al. Diabetes mellitus-Progress and opportunities in the evolving epidemic. Cell 187, 3789–3820 (2024). 10.1016/j.cell.2024.06.029

2 Chen, S., et al. The global macroeconomic burden of diabetes mellitus. Nat Med 32, 126–138 (2026). 10.1038/s41591-025-04027-5

3 Genitsaridi, I., et al. 11th edition of the IDF Diabetes Atlas: global, regional, and national diabetes prevalence estimates for 2024 and projections for 2050. Lancet Diabetes Endocrinol 14, 149–156 (2026). 10.1016/S2213-8587(25)00299-2

4 Collaboration, N. C. D. R. F. Worldwide trends in diabetes prevalence and treatment from 1990 to 2022: a pooled analysis of 1108 population-representative studies with 141 million participants. Lancet 404, 2077–2093 (2024). 10.1016/S0140-6736(24)02317-1

5 Lehrstrand, J., et al. Illuminating the complete ss-cell mass of the human pancreas-signifying a new view on the islets of Langerhans. Nat Commun 15, 3318 (2024). 10.1038/s41467-024-47686-7

6 Evans-Molina, C., et al. Heterogeneous endocrine cell composition defines human islet functional phenotypes. Nat Commun 17 (2026). 10.1038/s41467-026-70689-5

7 Klein, L., et al. Explainable AI-based analysis of human pancreas sections identifies traits of type 2 diabetes. Nat Commun 17, 1558 (2026). 10.1038/s41467-026-69295-2

8 Gottmann, P., et al. Heterogeneous Development of beta-Cell Populations in Diabetes-Resistant and-Susceptible Mice. Diabetes 71, 1962–1978 (2022). 10.2337/db21-1030

9 Wigger, L., et al. Multi-omics profiling of living human pancreatic islet donors reveals heterogeneous beta cell trajectories towards type 2 diabetes. Nat Metab 3, 1017–1031 (2021). 10.1038/s42255-021-00420-9

10 Butler, A. E., et al. Beta-cell deficit and increased beta-cell apoptosis in humans with type 2 diabetes. Diabetes 52, 102–110 (2003). 10.2337/diabetes.52.1.102

11 Moin, A. S. M. & Butler, A. E. Alterations in Beta Cell Identity in Type 1 and Type 2 Diabetes. Curr Diab Rep 19, 83 (2019). 10.1007/s11892-019-1194-6

12 Wang, Z., York, N. W., Nichols, C. G. & Remedi, M. S. Pancreatic beta cell dedifferentiation in diabetes and redifferentiation following insulin therapy. Cell Metab 19, 872–882 (2014). 10.1016/j.cmet.2014.03.010

13 Zhang, N., et al. Intrapancreatic adipocytes and beta cell dedifferentiation in human type 2 diabetes. Diabetologia 68, 1242–1260 (2025). 10.1007/s00125-025-06392-9

14 Narasimhan, A., Flores, R. R., Robbins, P. D. & Niedernhofer, L. J. Role of Cellular Senescence in Type II Diabetes. Endocrinology 162 (2021). 10.1210/endocr/bqab136

15 Talchai, C., Xuan, S., Lin, H. V., Sussel, L. & Accili, D. Pancreatic beta cell dedifferentiation as a mechanism of diabetic beta cell failure. Cell 150, 1223–1234 (2012). 10.1016/j.cell.2012.07.029

16 Brown, M. E., et al. Pancreatic islet beta-cell subtypes are derived from biochemically-distinct and nutritionally-regulated islet progenitors. Nat Commun 16, 5758 (2025). 10.1038/s41467-025-60831-0

17 Cohrs, C. M., et al. Dysfunction of Persisting beta Cells Is a Key Feature of Early Type 2 Diabetes Pathogenesis. Cell Rep 31, 107469 (2020). 10.1016/j.celrep.2020.03.033

18 Abdulreda, M. H., Rodriguez-Diaz, R., Caicedo, A. & Berggren, P. O. Liraglutide Compromises Pancreatic beta Cell Function in a Humanized Mouse Model. Cell Metab 23, 541–546 (2016). 10.1016/j.cmet.2016.01.009

19 van Raalte, D. H. & Verchere, C. B. Glucagon-Like Peptide-1 Receptor Agonists: Beta-Cell Protection or Exhaustion? Trends Endocrinol Metab 27, 442–445 (2016). 10.1016/j.tem.2016.04.009

20 Basile, G., Kulkarni, R. N. & Morgan, N. G. How, When, and Where Do Human beta-Cells Regenerate? Curr Diab Rep 19, 48 (2019). 10.1007/s11892-019-1176-8

21 Marquard, J., et al. Characterization of pancreatic NMDA receptors as possible drug targets for diabetes treatment. Nat Med 21, 363–372 (2015). 10.1038/nm.3822

22 Marquard, J., et al. Effects of dextromethorphan as add-on to sitagliptin on blood glucose and serum insulin concentrations in individuals with type 2 diabetes mellitus: a randomized, placebo-controlled, double-blinded, multiple crossover, single-dose clinical trial. Diabetes Obes Metab 18, 100–103 (2016). 10.1111/dom.12576

23 Dupuis, J. P., Nicole, O. & Groc, L. NMDA receptor functions in health and disease: Old actor, new dimensions. Neuron 111, 2312–2328 (2023). 10.1016/j.neuron.2023.05.002

24 Petersen, J., et al. GLP-1-directed NMDA receptor antagonism for obesity treatment. Nature 629, 1133–1141 (2024). 10.1038/s41586-024-07419-8

25 Sterk, M., et al. NMDA receptor inhibition increases, synchronizes, and stabilizes the collective pancreatic beta cell activity: Insights through multilayer network analysis. PLoS Comput Biol 17, e1009002 (2021). 10.1371/journal.pcbi.1009002

26 Welters, A., Lammert, E., Mayatepek, E. & Meissner, T. Need for Better Diabetes Treatment: The Therapeutic Potential of NMDA Receptor Antagonists. Klin Padiatr 229, 14–20 (2017). 10.1055/s-0042-117831

27 Welters, A., et al. NMDAR antagonists for the treatment of diabetes mellitus-Current status and future directions. Diabetes Obes Metab 19 Suppl 1, 95–106 (2017). 10.1111/dom.13017

28 Scholz, O., et al. Peripherally active dextromethorphan derivatives lower blood glucose levels by targeting pancreatic islets. Cell Chem Biol 28, 1474–1488 e1477 (2021). 10.1016/j.chembiol.2021.05.011

29 Pelligra, A., et al. Pancreatic islet protection at the expense of secretory function involves serine-linked mitochondrial one-carbon metabolism. Cell Rep 42, 112615 (2023). 10.1016/j.celrep.2023.112615

30 Scholz, O., et al. Protection of pancreatic islets from oxidative cell death by a peripherally-active morphinan with increased drug safety. Mol Metab 75, 101775 (2023). 10.1016/j.molmet.2023.101775

31 Roshanravan, H., Kim, E. Y. & Dryer, S. E. NMDA Receptors as Potential Therapeutic Targets in Diabetic Nephropathy: Increased Renal NMDA Receptor Subunit Expression in Akita Mice and Reduced Nephropathy Following Sustained Treatment With Memantine or MK-801. Diabetes 65, 3139–3150 (2016). 10.2337/db16-0209

32 Gresch, A., et al. Selective Inhibition of N-Methyl-d-aspartate Receptors with GluN2B Subunit Protects beta Cells against Stress-Induced Apoptotic Cell Death. J Pharmacol Exp Ther 379, 235–244 (2021). 10.1124/jpet.121.000807

33 Yagan, M., et al. Atf4 Protects Islet beta-Cell Identity and Function Under Acute Glucose-Induced Stress but Promotes beta-Cell Failure in the Presence of Free Fatty Acid. Diabetes 74, 838–849 (2025). 10.2337/db24-0360

34 Weippert, C., et al. CHAC1-linked NRF2 protein expression participates in pancreatic islet protection by dextrorphan. iScience 29 (2026). 10.1016/j.isci.2026.117474

35 Balakrishnan, A., et al. Analysis of ROR1 Protein Expression in Human Cancer and Normal Tissues. Clin Cancer Res 23, 3061–3071 (2017). 10.1158/1078-0432.CCR-16-2083

36 Kurita, Y., et al. A High-Fat/High-Sucrose Diet Induces WNT4 Expression in Mouse Pancreatic beta-cells. Kurume Med J 65, 55–62 (2019). 10.2739/kurumemedj.MS652008

37 Gerritsen, J. S. & White, F. M. Phosphoproteomics: a valuable tool for uncovering molecular signaling in cancer cells. Expert Rev Proteomics 18, 661–674 (2021). 10.1080/14789450.2021.1976152

38 Tang, J. S., Li, Q. R., Li, J. M., Wu, J. R. & Zeng, R. Systematic Synergy of Glucose and GLP-1 to Stimulate Insulin Secretion Revealed by Quantitative Phosphoproteomics. Sci Rep 7, 1018 (2017). 10.1038/s41598-017-00841-1

39 Yang, P., et al. KinasePA: Phosphoproteomics data annotation using hypothesis driven kinase perturbation analysis. Proteomics 16, 1868–1871 (2016). 10.1002/pmic.201600068

40 Kim, H. J., Kim, T., Xiao, D. & Yang, P. Protocol for the processing and downstream analysis of phosphoproteomic data with PhosR. STAR Protoc 2, 100585 (2021). 10.1016/j.xpro.2021.100585

41 Sacco, F., et al. Phosphoproteomics Reveals the GSK3-PDX1 Axis as a Key Pathogenic Signaling Node in Diabetic Islets. Cell Metab 29, 1422–1432 e1423 (2019). 10.1016/j.cmet.2019.02.012

42 Doliba, N. M., et al. Inhibition of cholinergic potentiation of insulin secretion from pancreatic islets by chronic elevation of glucose and fatty acids: Protection by casein kinase 2 inhibitor. Mol Metab 6, 1240–1253 (2017). 10.1016/j.molmet.2017.07.017

43 Fazakerley, D. J., et al. Phosphoproteomics reveals rewiring of the insulin signaling network and multi-nodal defects in insulin resistance. Nat Commun 14, 923 (2023). 10.1038/s41467-023-36549-2

44 Astapova, I., et al. The nuclear receptor corepressor (NCoR) controls thyroid hormone sensitivity and the set point of the hypothalamic-pituitary-thyroid axis. Mol Endocrinol 25, 212–224 (2011). 10.1210/me.2010-0462

45 Shimizu, H., et al. 7260 Nuclear Corepressor 1 (NCoR1) plays important role in the regulation of insulin gene transcription in mouse Min6 pancreatic beta cells. Journal of the Endocrine Society 8 (2024). 10.1210/jendso/bvae163.870

46 Walth-Hummel, A. A., et al. TBL1X/TBL1XR1 govern beta-cell identity through a PAX6-containing gene regulatory network. Nat Commun 17 (2026). 10.1038/s41467-026-72077-5

47 Remsberg, J. R., et al. Deletion of histone deacetylase 3 in adult beta cells improves glucose tolerance via increased insulin secretion. Mol Metab 6, 30–37 (2017). 10.1016/j.molmet.2016.11.007

48 Garofalo, R. S., et al. Severe diabetes, age-dependent loss of adipose tissue, and mild growth deficiency in mice lacking Akt2/PKB beta. J Clin Invest 112, 197–208 (2003). 10.1172/JCI16885

49 Haggblad Sahlberg, S., et al. Different functions of AKT1 and AKT2 in molecular pathways, cell migration and metabolism in colon cancer cells. Int J Oncol 50, 5–14 (2017). 10.3892/ijo.2016.3771

50 Manning, B. D. & Toker, A. AKT/PKB Signaling: Navigating the Network. Cell 169, 381–405 (2017). 10.1016/j.cell.2017.04.001

51 Leibiger, B., et al. PI3K-C2alpha Knockdown Results in Rerouting of Insulin Signaling and Pancreatic Beta Cell Proliferation. Cell Rep 13, 15–22 (2015). 10.1016/j.celrep.2015.08.058

52 Knoch, K. P., et al. PTBP1 is required for glucose-stimulated cap-independent translation of insulin granule proteins and Coxsackieviruses in beta cells. Mol Metab 3, 518–530 (2014). 10.1016/j.molmet.2014.05.002

53 Knoch, K. P., et al. cAMP-dependent phosphorylation of PTB1 promotes the expression of insulin secretory granule proteins in beta cells. Cell Metab 3, 123–134 (2006). 10.1016/j.cmet.2005.12.008

54 Lytrivi, M., Tong, Y., Virgilio, E., Yi, X. & Cnop, M. Diabetes mellitus and the key role of endoplasmic reticulum stress in pancreatic beta cells. Nat Rev Endocrinol 21, 546–563 (2025). 10.1038/s41574-025-01129-5

55 Aoto, M., et al. The splicing factor Acin1 is essential for embryonic development but has limited effects on muscle structure and homeostasis. Sci Rep 15, 14017 (2025). 10.1038/s41598-025-98851-x

56 Kelly, S., et al. Single-Islet Proteomics Maps Pseudo-Temporal Islet Immune Responses and Dysfunction in Stage 1 Type 1 Diabetes. bioRxiv (2025). 10.1101/2025.11.10.687674

57 Shibue, K., et al. Genome-wide CRISPR Screen Identifies Sec31A as a Key Regulator of Alpha Cell Survival. Nat Commun 16, 9159 (2025). 10.1038/s41467-025-64169-5

58 Su, K. H., et al. HSF1 critically attunes proteotoxic stress sensing by mTORC1 to combat stress and promote growth. Nat Cell Biol 18, 527–539 (2016). 10.1038/ncb3335

59 Li, W., et al. In situ structure of actin remodeling during glucose-stimulated insulin secretion using cryo-electron tomography. Nat Commun 15, 1311 (2024). 10.1038/s41467-024-45648-7

60 Leduc, M., et al. ERK1 is dispensable for mouse pancreatic beta cell function but is necessary for glucose-induced full activation of MSK1 and CREB. Diabetologia 60, 1999–2010 (2017). 10.1007/s00125-017-4356-6

61 Costes, S., et al. ERK1/2 control phosphorylation and protein level of cAMP-responsive element-binding protein: a key role in glucose-mediated pancreatic beta-cell survival. Diabetes 55, 2220–2230 (2006). 10.2337/db05-1618

62 Alessi, D. R., Cuenda, A., Cohen, P., Dudley, D. T. & Saltiel, A. R. PD 098059 is a specific inhibitor of the activation of mitogen-activated protein kinase kinase in vitro and in vivo. J Biol Chem 270, 27489–27494 (1995). 10.1074/jbc.270.46.27489

63 Dudley, D. T., Pang, L., Decker, S. J., Bridges, A. J. & Saltiel, A. R. A synthetic inhibitor of the mitogen-activated protein kinase cascade. Proc Natl Acad Sci U S A 92, 7686–7689 (1995). 10.1073/pnas.92.17.7686

64 Hrovatin, K., et al. Delineating mouse beta-cell identity during lifetime and in diabetes with a single cell atlas. Nat Metab 5, 1615–1637 (2023). 10.1038/s42255-023-00876-x

65 Golson, M. L., et al. Activated FoxM1 attenuates streptozotocin-mediated beta-cell death. Mol Endocrinol 28, 1435–1447 (2014). 10.1210/me.2014-1024

66 Kawamori, D., et al. Insulin signaling in alpha cells modulates glucagon secretion in vivo. Cell Metab 9, 350–361 (2009). 10.1016/j.cmet.2009.02.007

67 Chau, G. C., et al. mTOR controls ChREBP transcriptional activity and pancreatic beta cell survival under diabetic stress. J Cell Biol 216, 2091–2105 (2017). 10.1083/jcb.201701085

68 Gu, Y., Lindner, J., Kumar, A., Yuan, W. & Magnuson, M. A. Rictor/mTORC2 is essential for maintaining a balance between beta-cell proliferation and cell size. Diabetes 60, 827–837 (2011). 10.2337/db10-1194

69 Jaafar, R., et al. mTORC1 to AMPK switching underlies beta-cell metabolic plasticity during maturation and diabetes. J Clin Invest 129, 4124–4137 (2019). 10.1172/JCI127021

70 Brown, E. J., et al. A mammalian protein targeted by G1-arresting rapamycin-receptor complex. Nature 369, 756–758 (1994). 10.1038/369756a0

71 Schaffer, L., et al. A novel high-affinity peptide antagonist to the insulin receptor. Biochem Biophys Res Commun 376, 380–383 (2008). 10.1016/j.bbrc.2008.08.151

72 Okawa, E. R., et al. Essential roles of insulin and IGF-1 receptors during embryonic lineage development. Mol Metab 47, 101164 (2021). 10.1016/j.molmet.2021.101164

73 Iida, M., Harari, P. M., Wheeler, D. L. & Toulany, M. Targeting AKT/PKB to improve treatment outcomes for solid tumors. Mutat Res 819–820, 111690 (2020). 10.1016/j.mrfmmm.2020.111690

74 Sharma, K., et al. Ultradeep human phosphoproteome reveals a distinct regulatory nature of Tyr and Ser/Thr-based signaling. Cell Rep 8, 1583–1594 (2014). 10.1016/j.celrep.2014.07.036

75 Eberhard, D., et al. Semaphorin-3A regulates liver sinusoidal endothelial cell porosity and promotes hepatic steatosis. Nat Cardiovasc Res 3, 734–753 (2024). 10.1038/s44161-024-00487-z

76 Wong, E. H., et al. The anticonvulsant MK-801 is a potent N-methyl-D-aspartate antagonist. Proc Natl Acad Sci U S A 83, 7104–7108 (1986). 10.1073/pnas.83.18.7104

77 Konstantinova, I., et al. EphA-Ephrin-A-mediated beta cell communication regulates insulin secretion from pancreatic islets. Cell 129, 359–370 (2007). 10.1016/j.cell.2007.02.044

78 Jain, R., et al. Pharmacological inhibition of Eph receptors enhances glucose-stimulated insulin secretion from mouse and human pancreatic islets. Diabetologia 56, 1350–1355 (2013). 10.1007/s00125-013-2877-1

79 Volta, F., et al. Glucose homeostasis is regulated by pancreatic β-cell cilia via endosomal EphA-processing. Nat Commun 10, 5686 (2019). 10.1038/s41467-019-12953-5

80 Kulkarni, R. N. & Kahn, C. R. Ephs and ephrins keep pancreatic Beta cells connected. Cell 129, 241–243 (2007). 10.1016/j.cell.2007.04.006

81 Kipps, T. J. ROR1: an orphan becomes apparent. Blood 140, 1583–1591 (2022). 10.1182/blood.2021014760

82 Nomi, M., et al. Loss of mRor1 enhances the heart and skeletal abnormalities in mRor2-deficient mice: redundant and pleiotropic functions of mRor1 and mRor2 receptor tyrosine kinases. Mol Cell Biol 21, 8329–8335 (2001). 10.1128/MCB.21.24.8329-8335.2001

83 Yamaguchi, T., et al. ROR1 sustains caveolae and survival signalling as a scaffold of cavin-1 and caveolin-1. Nat Commun 7, 10060 (2016). 10.1038/ncomms10060

84 Lin, W., et al. IGFBP5 is an ROR1 ligand promoting glioblastoma invasion via ROR1/HER2-CREB signaling axis. Nat Commun 14, 1578 (2023). 10.1038/s41467-023-37306-1

85 Cerpa, W., Latorre-Esteves, E. & Barria, A. RoR2 functions as a noncanonical Wnt receptor that regulates NMDAR-mediated synaptic transmission. Proc Natl Acad Sci U S A 112, 4797–4802 (2015). 10.1073/pnas.1417053112

86 Coppieters, K. T., Wiberg, A., Amirian, N., Kay, T. W. & von Herrath, M. G. Persistent glucose transporter expression on pancreatic beta cells from longstanding type 1 diabetic individuals. Diabetes Metab Res Rev 27, 746–754 (2011). 10.1002/dmrr.1246

87 De Vos, A., et al. Human and rat beta cells differ in glucose transporter but not in glucokinase gene expression. J Clin Invest 96, 2489–2495 (1995). 10.1172/JCI118308

88 Reisberg, B., et al. Memantine in moderate-to-severe Alzheimer’s disease. N Engl J Med 348, 1333–1341 (2003). 10.1056/NEJMoa013128

89 Tu, W., et al. DAPK1 interaction with NMDA receptor NR2B subunits mediates brain damage in stroke. Cell 140, 222–234 (2010). 10.1016/j.cell.2009.12.055

90 Bader, E., et al. Identification of proliferative and mature beta-cells in the islets of Langerhans. Nature 535, 430–434 (2016). 10.1038/nature18624

91 Barlow, A. D., Nicholson, M. L. & Herbert, T. P. Evidence for rapamycin toxicity in pancreatic beta-cells and a review of the underlying molecular mechanisms. Diabetes 62, 2674–2682 (2013). 10.2337/db13-0106

92 Yuan, T., Lupse, B., Maedler, K. & Ardestani, A. mTORC2 Signaling: A Path for Pancreatic beta Cell’s Growth and Function. J Mol Biol 430, 904–918 (2018). 10.1016/j.jmb.2018.02.013

93 Gouda, M. M. Rheb1 as a novel beta-cell regulator connecting mTORC1, AMPK, and NOTCH1 pathways for efficient diabetes therapy. World J Diabetes 16, 108310 (2025). 10.4239/wjd.v16.i8.108310

94 Yang, Y., Song, W. J. & Zhang, J. J. Ras homolog enriched in brain 1 regulates beta cell mass and beta cell function via mTORC1/AMPK/Notch1 pathways. World J Diabetes 16, 104973 (2025). 10.4239/wjd.v16.i6.104973

95 Mafi, S., et al. The mTOR Signaling Pathway Interacts with the ER Stress Response and the Unfolded Protein Response in Cancer. Cancer Res 83, 2450–2460 (2023). 10.1158/0008-5472.CAN-22-3032

96 Szwed, A., Kim, E. & Jacinto, E. Regulation and metabolic functions of mTORC1 and mTORC2. Physiol Rev 101, 1371–1426 (2021). 10.1152/physrev.00026.2020

97 Asahara, S. I., Inoue, H., Watanabe, H. & Kido, Y. Roles of mTOR in the Regulation of Pancreatic beta-Cell Mass and Insulin Secretion. Biomolecules 12, 614 (2022). 10.3390/biom12050614

98 Li, R., Wilson, K. F. & Cerione, R. A. Elucidation of an mTORC2-PKC-NRF2 pathway that sustains the ATF4 stress response and identification of Sirt5 as a key ATF4 effector. Cell Death Discov 8, 357 (2022). 10.1038/s41420-022-01156-5

99 Hakonen, E., et al. In vivo activation of the PI3K-Akt pathway in mouse beta cells by the EGFR mutation L858R protects against diabetes. Diabetologia 57, 970–979 (2014). 10.1007/s00125-014-3175-2

100 Bernal-Mizrachi, E., et al. Defective insulin secretion and increased susceptibility to experimental diabetes are induced by reduced Akt activity in pancreatic islet beta cells. J Clin Invest 114, 928–936 (2004). 10.1172/JCI20016

101 Rao, P., et al. Gene transfer of constitutively active Akt markedly improves human islet transplant outcomes in diabetic severe combined immunodeficient mice. Diabetes 54, 1664–1675 (2005). 10.2337/diabetes.54.6.1664

102 Yoon, S. C., et al. The effect of MK-801 on mTOR/p70S6K and translation-related proteins in rat frontal cortex. Neurosci Lett 434, 23–28 (2008). 10.1016/j.neulet.2008.01.020

103 Petit-Pedrol, M. & Groc, L. Regulation of membrane NMDA receptors by dynamics and protein interactions. J Cell Biol 220 (2021). 10.1083/jcb.202006101

104 Trepanier, C. H., Jackson, M. F. & MacDonald, J. F. Regulation of NMDA receptors by the tyrosine kinase Fyn. FEBS J 279, 12–19 (2012). 10.1111/j.1742-4658.2011.08391.x

105 Grant, S. G. Analysis of NMDA receptor mediated synaptic plasticity using gene targeting: roles of Fyn and FAK non-receptor tyrosine kinases. J Physiol Paris 90, 337–338 (1996). 10.1016/s0928-4257(97)87914-4

106 Abe, T., et al. Fyn kinase-mediated phosphorylation of NMDA receptor NR2B subunit at Tyr1472 is essential for maintenance of neuropathic pain. Eur J Neurosci 22, 1445–1454 (2005). 10.1111/j.1460-9568.2005.04340.x

107 Salter, M. W. & Kalia, L. V. Src kinases: a hub for NMDA receptor regulation. Nat Rev Neurosci 5, 317–328 (2004). 10.1038/nrn1368

108 Karachaliou, N., et al. ROR1 as a novel therapeutic target for EGFR-mutant non-small-cell lung cancer patients with the EGFR T790M mutation. Transl Lung Cancer Res 3, 122–130 (2014). 10.3978/j.issn.2218-6751.2014.03.02

109 Borcherding, N., Kusner, D., Liu, G. H. & Zhang, W. ROR1, an embryonic protein with an emerging role in cancer biology. Protein Cell 5, 496–502 (2014). 10.1007/s13238-014-0059-7

110 Yamaguchi, T., et al. NKX2-1/TITF1/TTF-1-Induced ROR1 is required to sustain EGFR survival signaling in lung adenocarcinoma. Cancer Cell 21, 348–361 (2012). 10.1016/j.ccr.2012.02.008

111 Sanchez-Lopez, E., et al. NF-kappaB-p62-NRF2 survival signaling is associated with high ROR1 expression in chronic lymphocytic leukemia. Cell Death Differ 27, 2206–2216 (2020). 10.1038/s41418-020-0496-1

112 Sheetz, J. B., et al. Structural Insights into Pseudokinase Domains of Receptor Tyrosine Kinases. Mol Cell 79, 390–405 e397 (2020). 10.1016/j.molcel.2020.06.018

113 Cerpa, W., Gambrill, A., Inestrosa, N. C. & Barria, A. Regulation of NMDA-receptor synaptic transmission by Wnt signaling. J Neurosci 31, 9466–9471 (2011). 10.1523/JNEUROSCI.6311-10.2011

114 Riquelme, R., Li, L., Gambrill, A. & Barria, A. ROR2 homodimerization is sufficient to activate a neuronal Wnt/calcium signaling pathway. J Biol Chem 299, 105350 (2023). 10.1016/j.jbc.2023.105350

115 Hudecek, M., et al. The B-cell tumor-associated antigen ROR1 can be targeted with T cells modified to express a ROR1-specific chimeric antigen receptor. Blood 116, 4532–4541 (2010). 10.1182/blood-2010-05-283309

116 Perez-Riverol, Y., et al. The PRIDE database resources in 2022: a hub for mass spectrometry-based proteomics evidences. Nucleic Acids Res 50, D543–D552 (2022). 10.1093/nar/gkab1038

117 Yesil, P., et al. A new collagenase blend increases the number of islets isolated from mouse pancreas. Islets 1, 185–190 (2009). 10.4161/isl.1.3.9556

118 Wu, T., et al. clusterProfiler 4.0: A universal enrichment tool for interpreting omics data. Innovation (Camb) 2, 100141 (2021). 10.1016/j.xinn.2021.100141

119 Yu, G. enrichplot: Visualization of Functional Enrichment Result. R package version 1.26.6. (2025). 10.18129/B9.bioc.enrichplot

120 Carlson, M. org.Mm.eg.db: Genome wide annotation for Mouse. R package version 3.20.0. (2024).

121 Pagès, H. C., M; Falcon, S; Li, N. AnnotationDbi: Manipulation of SQLite-based annotations in Bioconductor. R package version 1.68.0. (2024). doi.org/10.18129/B9.bioc.AnnotationDbi

122 Szklarczyk, D., et al. The STRING database in 2025: protein networks with directionality of regulation. Nucleic Acids Res 53, D730–D737 (2025). 10.1093/nar/gkae1113

123 Griess, K., et al. Sphingolipid subtypes differentially control proinsulin processing and systemic glucose homeostasis. Nat Cell Biol 25, 20–29 (2023). 10.1038/s41556-022-01027-2

124 Schindelin, J., et al. Fiji: an open-source platform for biological-image analysis. Nat Methods 9, 676–682 (2012). 10.1038/nmeth.2019

125 Hornbeck, P. V. et al. PhosphoSitePlus, 2014: mutations, PTMs and recalibrations. Nucleic Acids Res 43, D512–520 (2015). 10.1093/nar/gku1267

126 Hilhorst, R., et al. Peptide microarrays for profiling of serine/threonine kinase activity of recombinant kinases and lysates of cells and tissue samples. Methods Mol Biol 977, 259–271 (2013). 10.1007/978-1-62703-284-1_21

127 Metz, K. S., et al. Coral: Clear and Customizable Visualization of Human Kinome Data. Cell Syst 7, 347–350 e341 (2018). 10.1016/j.cels.2018.07.001

